# Structural and biochemical characterisation of the N-Carbamoyl-β-Alanine Amidohydrolase from *Rhizobium radiobacter* MDC 8606

**DOI:** 10.1101/2023.05.04.538398

**Authors:** Ani Paloyan, Armen Sargsyan, Mariam D. Karapetyan, Artur Hambardzumyan, Sergey Kocharov, Henry Panosyan, Karine Dyukova, Marina Kinosyan, Anna Krüger, Cecilia Piergentili, Will A. Stanley, Arnaud Baslé, Jon Marles-Wright, Garabed Antranikian

**Author notes:** To whom correspondence should be addressed Jon Marles-Wright, Ani Paloyan.

## Abstract

N-Carbamoyl-β-Alanine Amidohydrolase (CβAA) constitute one of the most important groups of industrially relevant enzymes used in production of optically pure amino acids and derivatives. In this study, a N-carbamoyl-β-alanine amidohydrolase encoding gene from *Rhizobium radiobacter* MDC 8606 was cloned and overexpressed in *Escherichia coli*. The purified recombinant enzyme (RrCβAA) showed a specific activity of 14 U/mg using N-carbamoyl-β-alanine as a substrate with an optimum activity of 55°C at pH 8.0. In this work, we report also the first prokaryotic N-carbamoyl-β-alanine amidohydrolases structure at a resolution of 2.0 Å. A discontinuous catalytic domain and a dimerization domain attached through a flexible hinge region at the domain interface has been revealed. We have found that the ligand is interacting with a conserved glutamic acid (Glu131), histidine (H385) and arginine (Arg291) residues. Studies let us to explain the preference on the enzyme for linear carbamoyl substrates as large carbamoyl substrates cannot fit in the active site of the enzyme. This work envisages the use of RrCβAA from the *Rhizobium radiobacter* MDC 8606 for the industrial production of L-α-, L-β-, and L-γ – amino acids. The structural analysis provides new insights on enzyme–substrate interaction, which shed light on engineering of N-carbamoyl-β-alanine amidohydrolases for high catalytic activity and broad substrate specificity.

## Introduction

Optically pure L-amino acids find many industrial uses, where they are used as feed and food additives and as intermediates for pharmaceuticals, cosmetics, and pesticides [1]. While there are only 20 standard proteinogenic amino acids, hundreds of amino acids have been identified in nature, or have been chemically synthesized [2]. Although they are less abundant than their proteinogenic L-α-analogues, natural and synthetic L-β-, L-γ-, and L-δ-amino acids have found applications in the pharmaceutical industry such as diaminobutiric acid [3], as well as in different fields of biotechnology such as being used to investigate the structure and dynamics of proteins, to study protein interactions, and to modulate the activity of proteins in living cells [4]. β-Amino acids, have been used as building blocks of peptides, peptidomimetics, and many other physiologically active compounds [5]; for example, β-alanine is used as a dietary supplement, especially by athletes for its potential activity in the formation of the dipeptides anserine and carnosine [6], which may improve cerebral blood flow and verbal episodic memory [7]. Another example is the well-known γ-aminobutyric acid and its derivatives, which are widely used as health supplements [8]. δ-Amino acids are particularly valuable as chemical precursors, for example 5-aminovalerate is a C5 platform chemical used in the synthesis of δ-valerolactam [9], glutarate [10], and as a precursor for nylon fibres [11], and resins [12].

In the last decade, chemical synthesis of these amino acids has received considerable research attention, and several reviews on catalytic asymmetric synthesis strategies can be found [13]. From the biotechnological point of view, among the amino acid production technologies, the hydantoinase process is distinguished as a multienzyme and ecologically friendly process, which guarantees absolute stereospecificity in the production of amino acids [14]. With this method, the potential production of any optically pure amino acids from a wide spectrum of D-, L-5-monosubstituted hydantoins has proven to be viable [1, 15]. The method is widely used for the production of L- and D-amino acids by using L-N-(E.C. 3.5.1.87) or D-N-carbamoylase enzymes (E.C. 3.5.1.77), which convert N-carbamoyl-amino acids to their corresponding optically pure amino acids in the last stage of the hydantoinase process [16-17]. Characterization of prokaryotic N-Carbamoyl-β-Alanine amidohydrolase enzymes (NCβAAs, E.C. 3.5.1.6) has opened a new route for the hydantoinase process, suggesting that the enzyme, due to its broad substrate spectrum, can be used to obtain not only L-α-, but also L-β-, L-γ-, L-δ-amino acids [1], thus opening up new application opportunities for an old enzyme. NCβAA is also able to hydrolyse non-substituted substrate analogues in which the carboxyl group is replaced by a sulfonic or phosphonic acid group [18]; however, very little biochemical or structural information is available for this enzyme and only four enzymes of prokaryotic origin have been characterized to date [18–21]. N-Carbamoyl-β-alanine amidohydrolase, also known as β-alanine synthase/β-ureidopropionase, is the third enzyme participating in the degradation of uracil and thymine, which converts N-carbamoyl-β-alanine and 2-methyl-N-carbamoyl-β-alanine to β-alanine and 2-methyl-β-alanine, respectively [22]. The structure/function relationships for eukaryotic versions of these enzymes have been determined [23-24]. Despite the same function, the prokaryotic versions of these enzymes are structurally and functionally more closely related to the bacterial N-L-carbamoylases [18]. There are unpublished crystal structures of amidohydrolases from *Burkholderia* species in the PDB, and L-N-carbamoylase of Geobacillus stearothermophilus CECT43 which has only 36 % amino acid sequence identity to RrCβAA. In this study, we present crystal structure of the *Rhizobium radiobacter* MDC 8606 N-carbamoyl-β-alanine amidohydrolase and assess its activity profile at different experimental conditions and determine its activity against a range of substrates. Our findings illuminate key specificity features compared with L-N-carbamoylases, which show activity toward only N-carbamoyl-α-amino acids. Our findings highlight the utility of this enzyme for a range of industrially relevant biotransformations producing valuable amino-acid products.

## Results and discussion

### Analysis of the R. radiobacter MDC 8606 CβAA protein sequence

The gene encoding *R. radiobacter* MDC 8606 CβAA (Rr CβAA) was amplified from the DNA of a strain held in the Microbial Depository Centre (MDC) of the SPC Armbiotechnology NAS RA, Armenia. Analysis of the translated protein sequence confirms that this protein is a member of the carbamoyl-amidohydrolase family (EC 3.5.1.6) (**Supplementary Figure 1**) with between 20 - 97 % amino acid sequence identity with enzymes in this family with demonstrated amidohydrolase activity. Based on analysis of the sequence activity relationships in this family, and the high degree of amino acid conservation in functionally important sites between the RrCβAA and bacterial N-carbamoyl-β-alanine amidohydrolases, we propose this enzyme as a bacterial L-N-carbamoylase in the peptidase M20 family [1].

**Figure 1.**
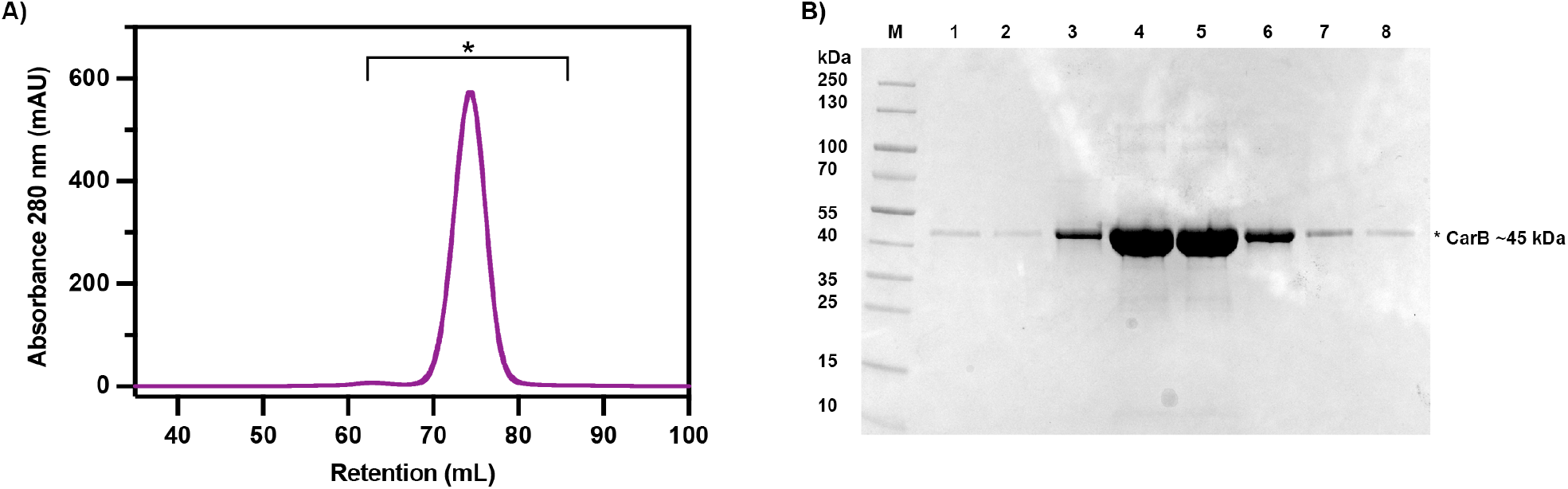
Purification of recombinant *RrCβAA*. A) Recombinant *RrCβAA* was purified by size exclusion chromatography after immobilised metal ion chromatography. The sample was run on a Superdex S200 16/60 column equilibrated with buffer containing 50 mM Tris.HCl pH 8.0, 150 mM NaCl. A single major peak at 74.4 mL is visible on the chromatogram. Peak fractions between 62 and 86 mL (labelled with a star) were collected for downstream analysis. B) SDS-PAGE of peak fractions (lanes 1-8) from the size exclusion chromatography run. The Fermentas pre-stained PageRuler was used at the molecular weight marker and the gel was stained with Coomassie brilliant blue stain.

### Production and purification of recombinant R. radiobacter MDC 8606 CβAA

To study the biochemical and structural properties of the RrCβAA protein, a plasmid was assembled to produce a C-terminally hexahistidine tagged recombinant version in *Escherichia coli* BL21(DE3). The protein was purified to homogeneity by a two-step purification procedure, using immobilised metal affinity chromatography (**Supplementary Figure 2**) and size exclusion chromatography (**Figure 1**). A single major peak was apparent on the size exclusion chromatogram at 74.4 mL, based on the calibration of this column, this can be ascribed to a protein with an apparent molecular weight of 90 kDa, this is consistent with the protein being a dimer in solution. **(Figure 1A)**. SDS-PAGE analysis shows a singlemain band of around 45 kDa, which is consistent with the calculated molecular weight of the protein of 44.7 kDa (**Figure 1B**).

**Figure 2.**
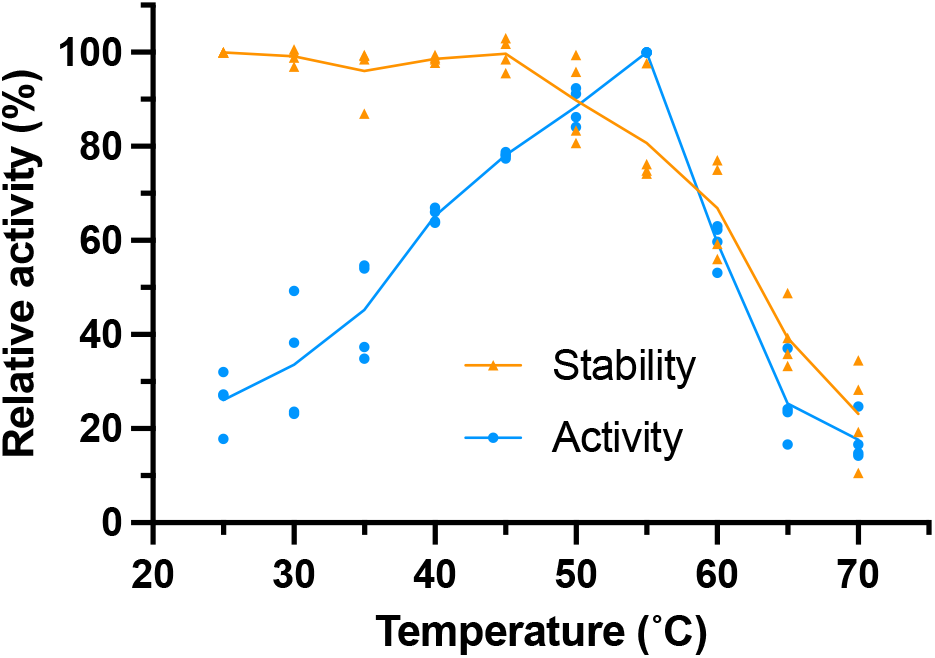
Activity and stability of RrCβAA with varying temperature. The activity and stability of the recombinant RrCβAA enzyme was assessed between 25 and 70°C. Experiments were performed with two technical replicates each from two biological replicates. Blue points show the activity profile over the temperature range at a fifteen-minute end point; the blue line represents the mean of the four measured replicates. Orange points show residual activity of enzyme after fifteen minutes pre-incubation over the temperature range prior to assay for fifteen minutes at 40°C; the orange line represents the mean of the four measured replicates.

The purified protein from the 74.4 mL size exclusion fraction was 30-times more active against N-carbamoyl β-alanine than the crude lysate and displayed an activity of around 13.4 U/mg under our standard assay conditions with the N-carbamoyl-L-β-alanine substrate (**Supplementary Table 1**).

**Table 1.** Comparison of biochemical properties of L-N-carbamoylases with respect to N-Carbamoyl-β-Alanine Amidohydrolases from *Rhizobium radiobacter* MDC 8606. For activity with metal cofactors, blue shading represents activation and orange inhibition, blank cells show data not available.

| Source | Specificity with Ureidopropionase substrate | Sub unit masses (kDa ) | Oligomer | Activity with metals |  |  |  |  |  |  |  |  |  | pH optimum | Temperature optimum (°C) | Ref |
| --- | --- | --- | --- | --- | --- | --- | --- | --- | --- | --- | --- | --- | --- | --- | --- | --- |
|  |  |  |  | C a | M n | F e | C o | N i | C u | Z n | A g | C d | H g |  |  |  |
| *R. radiobacter* MDC 8606 | Yes | 45.0 | monomer |  |  |  |  |  |  |  |  |  |  | ND | 55 | This study |
| *B. reuszeri* HSN1 | No | 44.3 | trimer |  |  |  |  |  |  |  |  |  |  | 8.5 | 50 | [35] |
| *A. tumefaciens* C58 (β-up) | Yes | 45.0 | dimer |  |  |  |  |  |  |  |  |  |  | 8 | 30 | [18] |
| *A. xylosoxidans* AKU 990 | ND | 65.0 | dimer |  |  |  |  |  |  |  |  |  |  | 8 - 8.3 | 30 | [36] |
| *A. aurescens* DSM3747 | No | 44.0 | dimer |  |  |  |  |  |  |  |  |  |  | 8.5 | 50 | [31] |
| *B. kaustophilus* CCRC1123 | No | 45.0 | ND |  |  |  |  |  |  |  |  |  |  | 7.4 | 70 | [37] |
| *P. putida* IFO 12996 (β-up) | Yes | 45.0 | dimer |  |  |  |  |  |  |  |  |  |  | 7.5-8.2 | 60 | [20] |
| *Pseudomonas* sp. NS671 | No | 45.0 | dimer |  |  |  |  |  |  |  |  |  |  | 7.5 | 40 | [32] |
| *B. stearothermophilus* NS1122A | ND | ND | ND |  |  |  |  |  |  |  |  |  |  | 8 | 60-70 | [33] |
| *G. stearothermophilus* CECT43 | ND | ND | ND |  |  |  |  |  |  |  |  |  |  | 7.5 | 65 | [34] |
| *Pseudomonas* sp. ON-4A | ND | 45.0 | tetramer |  |  |  |  |  |  |  |  |  |  | 9 | 50 | [30] |
| *S. meliloti* CECT4114 | No | 42.0 | dimer |  |  |  |  |  |  |  |  |  |  | 8.0 | 60 | [26] |

### R. radiobacter CβAA displays optimal activity between 50 and 60°C

To assess the impact of temperature on the activity profile of RrCβAA, the purified enzyme was assayed at temperatures between 25 and 70 °C for reaction times of fifteen minutes. The normalised reaction progress data shows a temperature optimum of 55 °C for the enzyme in the conditions tested (**Figure 2 blue line and Supplementary Table 2**). The thermostability of the enzyme was determined by incubating RrCβAA at different temperatures for fifteen minutes and assessing the residual activity at 40 °C. The enzyme displayed no significant reduction in activity up to 40 °C, with 50 % of activity lost at 65 °C (**Figure 2 orange line and Supplementary Table 3**).

### Divalent cations are required for Rr CβAA activity

The activity of the peptidase M20/M25/M40 family is known to be dependent on the presence of divalent cations in the active site to activate a catalytic water [25]. The purified RrCβAA enzyme was incubated in phosphate buffer with the addition of various cations, chelators, and reducing agents (**Figure 3A and Supplementary Table 4**). The addition of EDTA abolishes almost all enzyme activity, which is consistent with the requirement of metal cations for enzyme activity. The enzyme showed only 5 % activity after 1 hour incubation in the presence of 2 mM EDTA, whereas a continued overnight incubation fully inactivated RrCβAA. Assay of the prepared protein with the addition of divalent cations showed that Cd^2+^, Co^2+^, Ni2+, and Mg^2+^ had a strong positive effect on the enzyme activity, while Zn^2+^ and Cu^2+^ have distinct inhibitory effects on the purified enzyme.

**Figure 3.**
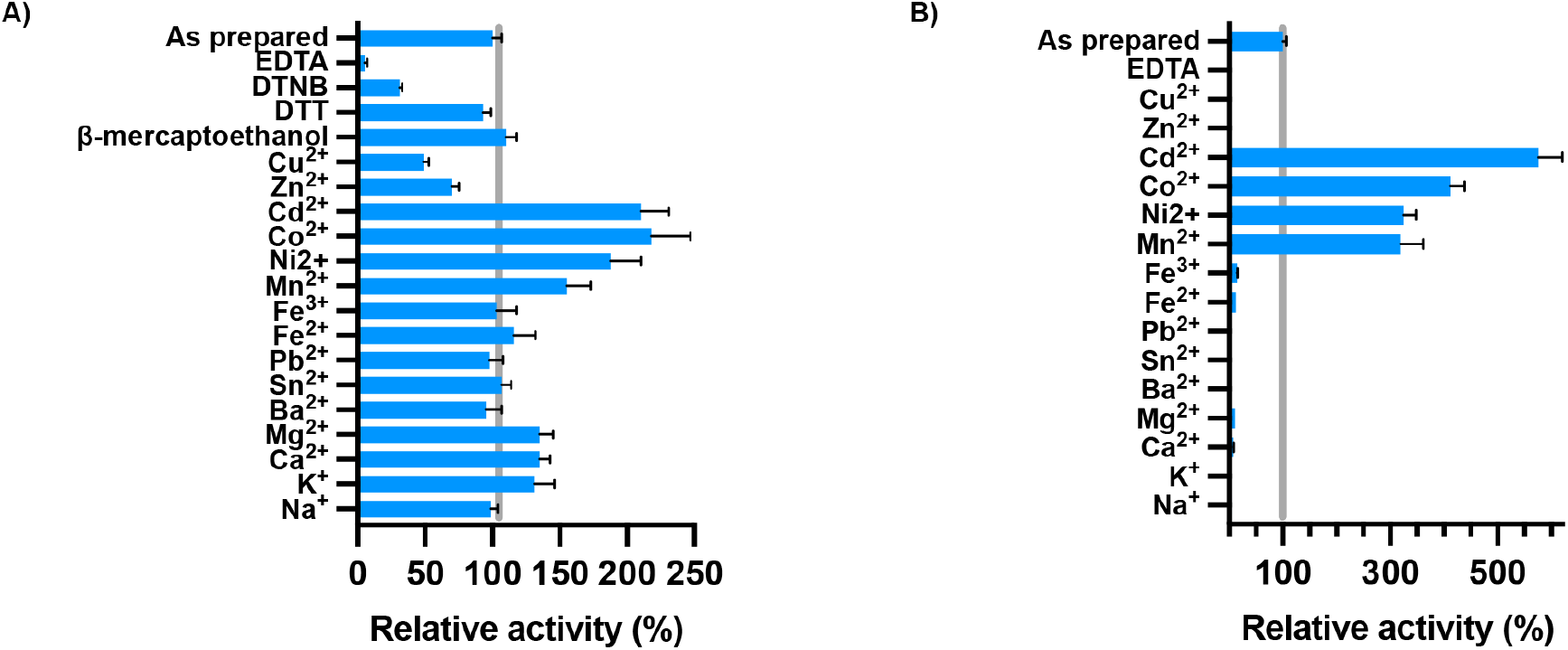
Effect of cations and chemical compounds on the activity of RrCβAA. A) The activity of the recombinant RrCβAA enzyme was assessed after 1 hour incubation at 4°C in the presence of 2 mM of different metals and EDTA, or 5 mM of DTNB, DTT, and β-mercaptoethanol. B) The recombinant RrCβAA enzyme was incubated with EDTA prior to the addition of different metals. Experiments were performed with three technical replicates each from two biological replicates. A specific activity of 13.6 U/mg obtained without additives was defined as 100 % activity.

In our experiments, full activity recovery of EDTA inactivated enzyme, was detected after incubation for one hour at 4 °C in phosphate buffer containing Mn^2+^, Ni^2+^, Co^2+^, or Cd^2+^, at 2 mM concentration (**Figure 3B and Supplementary Table 5**). While eukaryotic N-Carbamoyl-β-alanine amidohydrolase has been described as a Zn^2+^ dependent enzyme, our results show that the EDTA-inactivated RrCβAA is not recovered with Zn^2+^. Moreover, it shows an inhibitory effect on the purified recombinant enzyme, whereas Cu^2+^ was shown to be a stronger inhibitor for the RrCβAA enzyme. Enzyme activity was not affected by Fe^2+^, which is known as L-N-carbamoylase activator [26], nor by Sn^2+^ and Pb^2+^, known as inhibitors of *P. putida* IF0 12996 β-ureidopropionase [20]. Similar results were seen for βcar_At_ from *Agrobacterium tumefaciens* C58 [18], where the enzyme activity can be recovered with Mn^2+^, Ni^2+^, and Co^2+^. Interestingly the activity βcar_At_ could not be recovered with Cd^2+^, which is one of the preferred metal cations for RrCβAA. Moreover, Cd^2+^ shows an inhibitory effect on β-ureidopropionase from *Pseudomonas putida* IFO 12996 [20].

It is not possible to distinguish the physiological metal cation from these experiments, in the *E. coli* cytosol the recombinant enzyme is likely to be loaded with Zn^2+^; however, the metal binding site is clearly labile, as the cation is able to be removed with EDTA treatment and replaced with those added in excess in vitro.

Disulfide reducing agents such as β-mercaptoethanol and DTT do not show any inhibitory effects on the activity of the enzyme. Enzyme activity was not altered in the presence of 2 and 5 mM β-mercaptoethanol. Interestingly, the sulfhydryl reagent DTNB showed an inactivating effect on the enzyme. These results indicate that while cysteine residues do not play a key role in the enzyme activity, DTNB may form a covalent adduct that interferes with the activity of the enzyme through interactions with cysteine residues close to the active site. βcar_At_ from *Agrobacterium tumefaciens*

C58 was not inhibited by DTNB [18], whereas this compound showed inhibitory effect on other β-ureidopropionase and L-N-carbamoylase proteins.

### RrCβAA displays a broad substrate range with optimal activity against N-carbamoyl-L-β-alanine

To better understand the substrate specificity of the RrCβAA enzyme, we investigated its activity against N-carbamoyl-L-, D- and DL-amino acids (**Figure 4 and Supplementary Table 6**). The enzyme displays no activity toward N-carbamoyl-D-amino acids, with a clear stereo specificity toward N-carbamoyl-L-amino acids. Since the enzyme catalyses the third step in the pyrimidine degradation pathway, it shows greatest catalytic efficiency for N-carbamoyl-L-β-alanine. This is in contrast with the recombinant Atβcar from *Agrobacterium tumefaciens* C58, which displayed the highest activity toward N-carbamoyl-L-methionine [27]. The RrCβAA enzyme displayed a specific activity toward N-carbamoyl-L-α-alanine which was 2-fold lower than to N-carbamoyl-L-β-alanine, which differs only by the position of the carbamide group. In case of N-carbamoyl-L-β-alanine, the carbamide group is located at the edge of the β-carbon position, resulting in a linear structure. Similar results have been obtained with N-carbamoyl-α-amino- and N-carbamoyl-γ-amino butyric acids. The movement of the carbamide group from α-to the γ-position resulted in a more than 1.5-fold increase in the specific activity. Conversely, RrCβAA displayed very low activity against N-carbamoyl-L-valine and N-carbamoyl-L-leucine. Similar results were obtained with N-carbamoyl-L-α-phenyl-β-alanine and N-carbamoyl-L-β-phenyl-β-alanine. RrCβAA displayed good activity toward N-carbamoyl-L-methionine, which implies that the sulfur containing sidechain is accommodated within the active site in some way.

**Figure 4.**
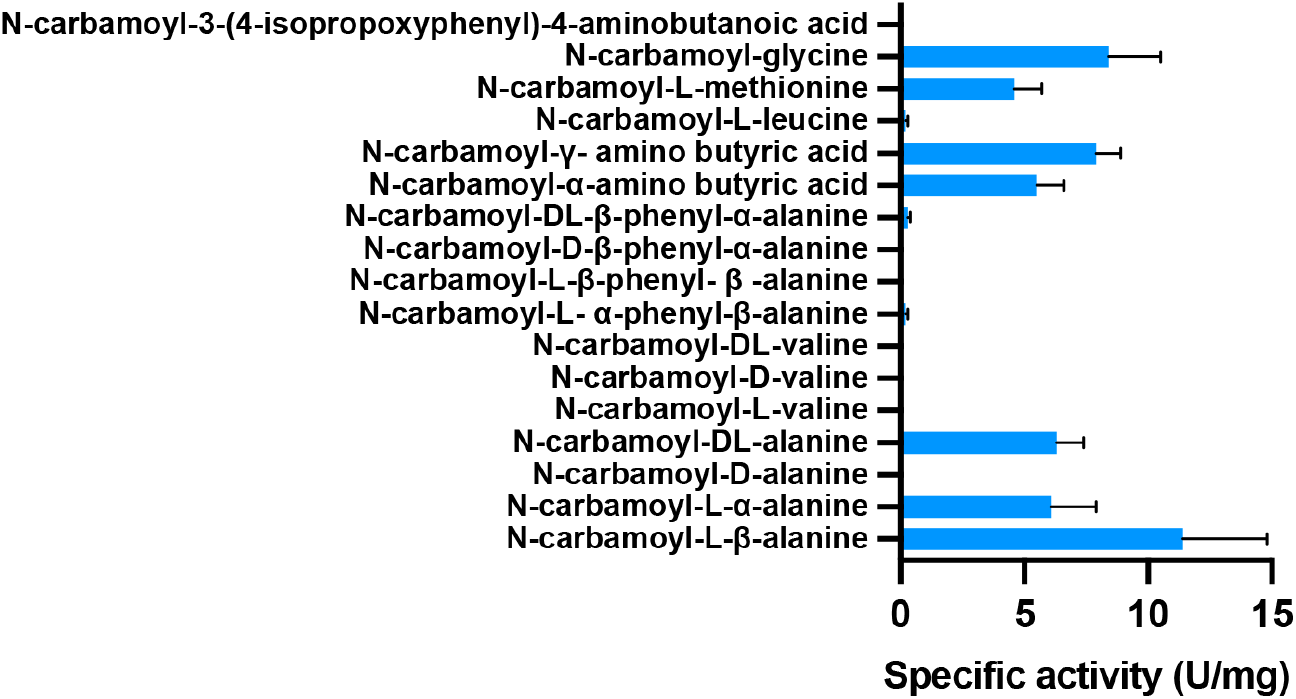
Substrate specificity of RrCβAA. The specific activity of purified RrCβAA was assessed toward various N-carbamoyl-amino acids. A reaction mixture containing 100 mM of different N-carbamoyl-amino acids was incubated at 40 °C for 10 min. The reaction was started by adding enzyme and was carried out for 15 min at 40 °C, pH 8.0. Experiments were performed with two technical replicates each from two biological replicates.

Our results indicate that RrCβAA has a distinct preference for N-carbamoyl-L-amino acids with linear R-groups and that its active site does not readily accommodate to the branched hydrophobic or aromatic sidechains.

### Crystal structure RrCβAA

Based on our results demonstrating a requirement for metal binding for catalysis and our exploration of the substrate preference of the RrCβAA, we determined crystal structure of RrCβAA to better understand the structure/function relationships that mediate substrate specificity in this enzyme. The structure of RrCβAA was determined to 2 Å resolution in P22121 space group, with two molecules in the asymmetric unit representing the functional dimer of the protein (**Supplementary Table 7**) (**Figure 5A**). The RrCβAA enzyme has a discontinuous catalytic domain from the N-terminus to residue 213 and residue 331 to the C-terminus. The dimerization domain is intercalated within the catalytic domain and comprises residues 214-330. The dimerization interface is formed between beta strands on one face of the domain (residues 269-278) and alpha helices on the opposite face (residues 230-260). The interface has a hydrophobic core, formed by the side chains of residues from both the strands and helices, a network of hydrogen bonds stabilising the beta-strand interface, and salt bridges across the top face of the alpha-helical interface (**Supplementary Figure 3**).

**Figure 5.**
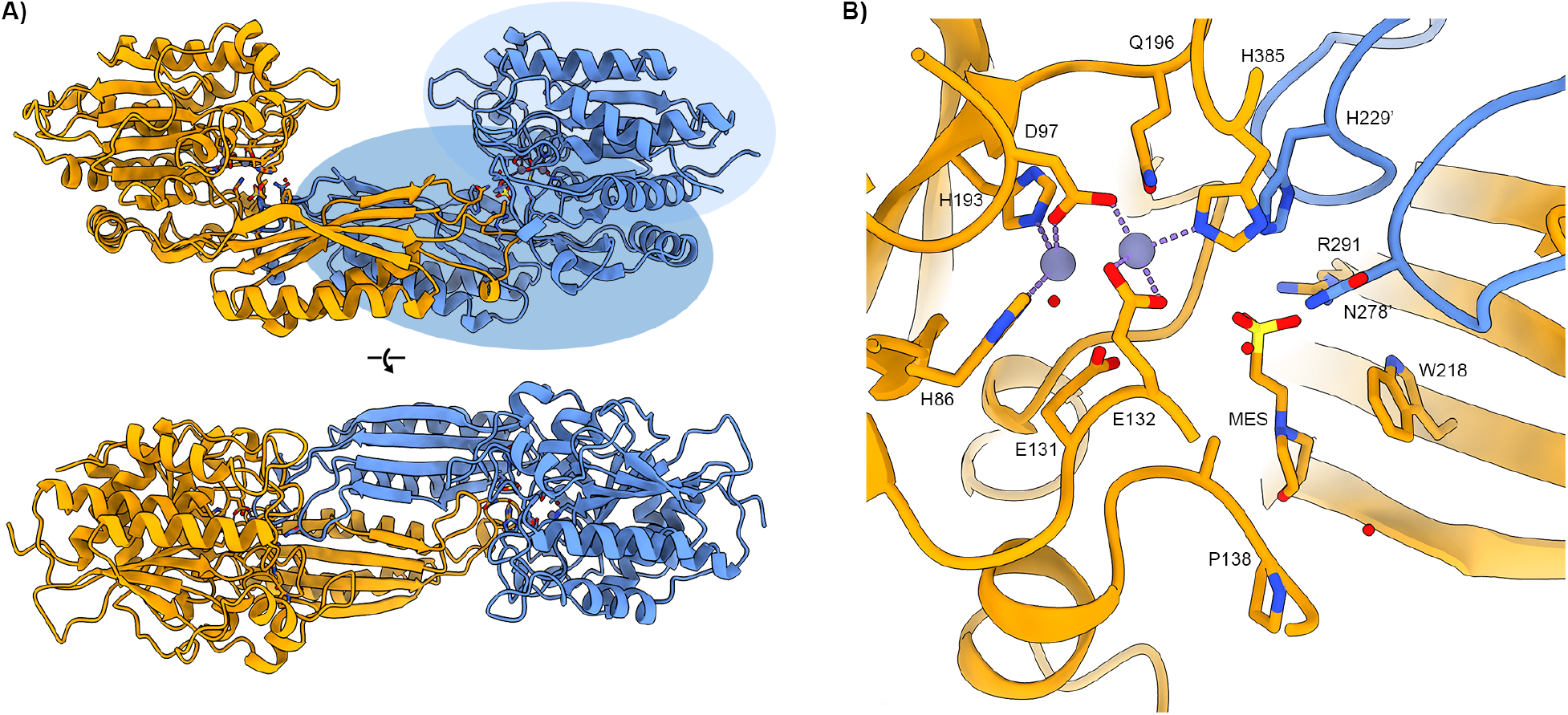
Crystal structure of RrCβAA. A) Overall structure of RrCβAA shown in cartoon depiction, monomers are coloured orange and blue. The dimerization domain of one monomer is highlighted in mid-blue, with the catalytic domain shown in light blue. B) Metal and ligand binding site of RrCβAA with interacting residues shown in stick representation coloured by atom. The ligand binding site comprises residues from both monomers, shown in orange and blue. Zinc ions are shown as purple spheres with coordinating bonds shown as purple dashes. A competing MES buffer ligand molecule from the crystallization condition is bound in the ligand binding site.

The catalytic and dimerization domains of RrCβAA are attached through a flexible hinge region at the domain interface. In other structural models of proteins in this family the catalytic domain rotates around this hinge to open and close the active site cleft. When aligned to other models, it is apparent that our RrCβAA model is in a partially closed state, with the catalytic domains of both chains in the asymmetric unit adopting essentially identical conformations. The hinging movement shown in previously determined models is found on a continuum between fully closed [28] and a wide-open state [29] (**Supplementary Figure 4A**). The models show a rotation range of approximately 45°; and when the catalytic domains are aligned, there is a relative 30 Å movement around the axis of rotation between the closed and open states at the end of the dimerization domain (**Supplementary Figure 4B**).

Each protein chain in the dimer has electron density features consistent with the presence of divalent cations in the putative metal binding site. These were modelled as Zn^2+^ ions based on the availability of zinc ions in *E. coli* expression host, and the presence of zinc in other published structures in this family. The Zn^2+^ ions coordinate conserved glutamic acid and histidine residues with ligand coordination distances of approximately 2.1 Å (**Figure 5B**). A strong peak of electron density was observed in the vicinity of the active site and a MES buffer molecule from the crystallisation condition was modelled in this region. The modelled MES refined well with good electron density fit and B-factors (**Figure 5B and Supplementary Figure 5**).

The presence of high concentrations of the competing MES buffer in the crystallisation condition hindered experiments to soak ligands, such as N-carbamoyl-beta-alanine, into the active site of the crystals to determine a structure of an enzyme ligand complex. Structural alignments of our RrCβAA model with other structures in this family with ligands in their active sites gives some insight into the ligand binding site (**Figure 6A**).

**Figure 6.**
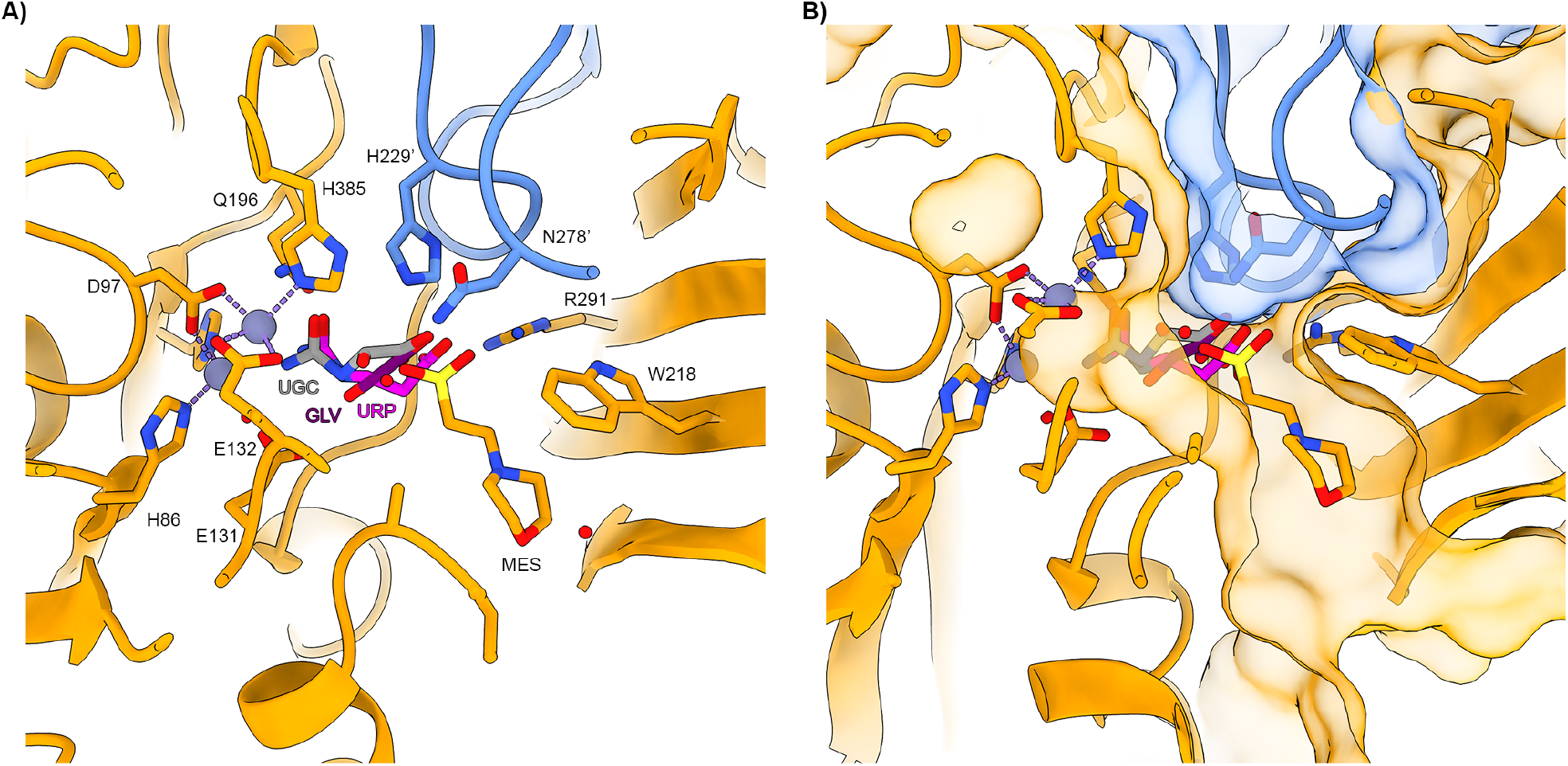
Productive ligand binding in RrCβAA is constrained by a tight active site cleft in the closed conformation. A) Structural homologues with bound ligands were aligned to the RrCβAA structural model. RrCβAA is shown as orange and blue cartoons with metal and ligand binding residues shown as stick representations with bound MES buffer shown. Modelled ligands are as follows: UGC – (S)-ureidoglycolate from PDB: 4PXB; GLV – beta-alanine from PDB 2V8G; URP – N-carbamoyl-beta-alanine from PDB: 5THW. B) Active site cleft shown with transparent surface to highlight the physical constraints placed on ligand binding in this space.

The carbamoyl group of the modelled ligands occupy a space close to the metal binding site where the group is oriented through interactions with the cations and cluster of conserved amino acids including glutamine (Gln196), glutamic acid (Glu131) and histidine residues (His385). The carboxylic acid group of the ligand forms a salt bridge with the conserved arginine residue (Arg291). These interactions essentially constrain the ligand binding at both functional groups. To form a productive ligand complex, the protein must engage ligand while in an open state and close around it to facilitate catalysis [29]. The observed preference for linear, and gamma substituted carbamoyl amino acids is a consequence of the steric constraints posed by the amino acids lining the active site cleft (**Figure 6B**). Large aromatic carbamoyl amino acids cannot fit within the closed active site and therefore the enzyme is not active against them. However, linear side chains at the alpha and beta positions may be accommodated in the active site cleft to form productive complexes.

## Conclusion

In this work we have demonstrated the recombinant production and activity of the *R. radiobacter* N-Carbamoyl-β-Alanine amidohydrolase enzyme (RrCβAA). RrCβAA was purified as a homodimer, like other β-ureidopropionase and L-N-carbamoylases, except L-N-carbamoylases characterize from *Brevibacillus reuszeri* HSN1 and *Pseudomonas sp*. ON-4 which have been shown to exist as a homotrimer and homotetramer, respectively [30]. Among studied carbamoylases with L-steriospecificity only N-Carbamoyl-β-Alanine amidohydrolase of *A. tumefaciens* C58 and L-N-carbamoylase of *P. putida* IFO 12996 have been demonstrated to have β-ureidopropionase activity (**Table 1**).

All β-ureidopropionase and L-N-carbamoylases are described as metalloenzymes and RrCβAA is no exception to this rule. The chelating agent EDTA abolishes the enzyme activity, which was recovered by the addition of Mn^2+^, Ni^2+^, Co^2+^, Cd^2+^. The first three metals are well known cofactors for this enzyme family; while Cd^2+^ has not been widely demonstrated as a cofactor for this enzyme, and in some cases has been shown to be inhibitory [20]. This is the first result showing that Cd^2+^ acts as a cofactor for this class of enzymes, although it is not clear from our structural analysis what the basis for these differences are. The other divalent cations tested, such as Cu^2+^, Zn^2+^ show an inhibitory effect on enzyme activity. These results indicate that the properties of RrCβAA are comparable to those of all known L-carbamoylases and β-ureidopropionase enzymes from other bacterial strains (**Table 1**).

Reducing compounds did not show an inhibitory effect on enzyme activity, this is consistent with our structural observations showing that there are no key cysteine residues involved in the catalysis. The enzyme is also not stabilised by any key disulphide bridges, which may be disrupted by reducing agents. The inhibitory effect of DTNB on the enzyme can be rationalised if it forms a covalent adduct with Cys364, which is close to the hinge region of the protein; such an adduct would prevent closure of the active site and inhibit the production of a catalytically competent intermediate state with any substrate.

The optimum activity for the RrCβAA was recorded at 55 °C, which is higher than the enzymes from *Agrobacterium tumefaciens* C58 [27], *Arthrobacter. aurescens* DSM3747 [31]. *Achromobacter xylosoxidans [20] Pseudomonas sp*. NS671 [32] and slightly lower than the enzymes from *P. putida* IFO12996 [20], *Bacillus stearothermophilus* NSl122A [33] and *Geobacillus stearothermophilus* CECT43 [34].

In terms of substrate preference and promiscuity the RrCβAA shows good activity against L-α-, L-β-, L-γ-amino acids, this contrasts with other L-carbamoylases described so far which show preferential activity to only N-carbamoyl L-α-amino acids. The lack of activity towards branched chain and aromatic amino acids limits its use against these substrates; however, there is certainly scope for employing focused mutagenesis to open the substrate binding site to accept these substrates. This strategy has been as demonstrated for the *S. meliloti* carbamoylase, which has been engineered to accept aromatic amino acids [28]. Further work on the RrCβAA enzyme will focus on expanding its substrate scope against these aromatic amino acids with high potential for use in industrially useful chemo-enzymatic cascades.

## Methods

### Reagents and substrates

Phusion® DNA Polymerase, BsaI restriction enzyme and T4 DNA ligase, were purchased from New England Biolabs (Hitchin, UK). Isopropyl β-D-1-thiogalactopyranoside (IPTG) was purchased from Merck, UK. The molecular weight marker for SDS–PAGE was purchased from Thermo Fisher Scientific (Cramlington, UK). Standards, and some substrates (N-carbamoyl-β-alanine (3-ureidopropionic acid), N-carbamoyl-glycine) were purchased from Sigma. Other N-carbamoyl-DL, L and D-amino acids have been synthesized for this study. ^1^H and ^13^C NMR analyses were performed to confirm their structures (**Supplementary Figure 6**). All other chemicals were of analytical grade.

### Bacterial strains and plasmids

The *Rhizobium radiobacter* MDC 8606 strain used as a source for the N-Carbamoyl-β-alanine amidohydrolase (RrCβAA) gene was taken from the Microbial Depository Center (MDC) of SPC “Armbiotechnology” NAS RA. *Escherichia coli* Top 10 and *E. coli* BL21 (DE3) strains were used for propagation of plasmids and protein expression respectively. A modified pET28 plasmid for Golden Gate cloning was a gift of Dr Laura Tuck.

### Nucleotide and amino acid sequence analysis

Sequence analysis of the RrCβAA gene was performed using the BLAST program [38]. Protein sequence alignments were performed in Multalin [39] and figures prepared with ESPript [40]. The nucleotide sequence data of the isolated RrCβAA, as well as 16s rRNA genes of *Rhizobium radiobacter* MDC 8606 strain were deposited in NCBI GeneBank database with the accession numbers MT542139 and MT534525.1 respectively.

### Cloning, expression, and purification of RrCβAA

To amplify the *R. radiobacter* MDC 8606 N-carbamoyl-β-alanine amidohydrolase open reading frame, primers RrCβAA-F (5’**GAC**GGTCTC**TA**ATGACGGCGGGTAAAAACTTGAC3’) and RrCβAA-R (5’**GAC**GGTCTC**TACCT**TTGCACGATCTCCGCAGTCTC3’) were designed using

*Agrobacterium tumefaciens* C58 N-carbamoyl-β-alanine amidohydrolase gene sequence as a template (GenBank: EF507843.1). PCR was performed using these primers against the purified *R. radiobacter* MDC 8606 genomic DNA with the following conditions: 98 °C for 1 min, followed by 30 cycles of 98 °C for 30 s, 60 °C for 30 s and 72 °C for 1 min, followed by a final elongation at 72 °C for 10 min. After examination by 1 % agarose TAE electrophoresis, the amplified product was purified by QIAquick PCR Purification Kit. The purified DNA fragment was then assembled via one-pot Golden Gate cloning [41] into a CIDAR MoClo [42] compatible pET28 vector via BsaI restriction sites introduced into the PCR product and pET28 vector. The resulting ligation product was transformed into chemically competent *E. coli* TOP10 cells with selection on LB agar plates supplemented with 35 μg/mL kanamycin, 1 mM IPTG, and 20 μg/mL X-Gal. Recombinant plasmid was extracted from white insert-positive clones by miniprep using a Qiagen Miniprep kit. The insert presence was confirmed by Sanger sequencing of the purified plasmids. The sequence verified plasmid was transformed into *E. coli* BL21(DE3) cells with selection on LB agar supplemented with 35 μg/mL kanamycin. A single colony was grown overnight at 37 °C in 100 mL LB medium, supplemented with 35 μg/mL kanamycin, with shaking at 180 rpm. The cells were sub-cultured into 2 L of LB, grown until OD_600_ 0.5, and recombinant protein production was induced with 1 mM IPTG, at 25°C, followed by incubation for a further 16 hours.

Cells were harvested by centrifugation at 7,000 × g for 20 min. The harvested cells were resuspended in 10 x w/v Buffer HisA (50 mM imidazole, 500 mM NaCl, 50 mM Tris-HCl, pH 8.0) and subsequently sonicated on ice for 5 minutes with 30 s on/off cycles at 60 watts power output. The lysate was cleared by centrifugation at 35,000 × g and filtered with a 0.45 μm syringe filter.

Cell free extract was applied to a 5 ml HisTrap FF column (GE Healthcare), and unbound proteins were washed off with 10 column volumes of Buffer HisA (50 mM Tris-HCl, pH 8.0, 500 mM NaCl, 50 mM imidazole). A step-gradient of 50 % and 100 % Buffer HisB (50 mM Tris-HCl, pH 8.0, 500 mM NaCl, 500 mM imidazole) was used to elute His-tagged proteins. Fractions of His-trap eluent containing the protein of interest (**Supplementary Figure 2**), were pooled, and concentrated by Vivaspin Turbo (Sartorius, 10 kDa MWCO) centrifugation devices at 4,000 × g, 18°C. The concentrated protein was then subjected to size-exclusion chromatography using an S200 16/60 column (Cytiva), equilibrated with Buffer GF (50 mM Tris-HCl, pH 8.0, 150 mM NaCl). Calibration data for the S200 16/60 gel filtration column used are available at https://doi.org/10.6084/m9.figshare.7752320.v1. Fractions were analysed by sodium dodecyl sulphate polyacrylamide gel electrophoresis using Mini-PROTEAN TGX precast 4-20% gels (BioRad) according to the standard method [43] to determine the molecular weight and the purity of the samples. Purified RrCβAA was concentrated and analysed by SDS–PAGE, after incubating the sample for 5 min at 95°C temperature, in the presence of 5 mM β-mercaptoethanol. The Fermentas pre-stained PageRuler was used as a protein molecular weight marker for SDS-PAGE. Gels were stained with Coomassie brilliant blue for visualisation of protein bands.

For characterization studies, purified RrCβAA was placed into 100 mM phosphate buffer, pH 8.0 (Tris-HCl shows absorption in the presence of the ortho-phthalaldehyde reagent) and stored at −80°C with the addition of 50 % (v/v) glycerol for enzyme characterization.

### General procedure for synthesis of carbamoyl amino acids

All chemicals used for synthesis were of analytical or reagent grade. N-Carbamoyl-β-Ala (**15**) was from “Sigma”. The compounds 2 – 14 were prepared using the amino acids from Reanal (Budapest, Hungary). Melting points were determined on a Boetius PHMK 76/0904 hot stage microscope (GDR) and are uncorrected. ^1^H and ^13^C NMR spectra were recorded on a Varian

Mercury-300 spectrometer, operating at 300 MHz; chemical shifts are reported in *δ* values (ppm) relative to tetramethylsilane as internal standard. Coupling constants (*J* values) are given in Hertz (Hz). The solvents mixture was DMSO-d_6_/CCl_4_, NMR spectra and assignments are shown in the Supplementary Information (**Supplementary Figure 6**); the signals are reported as follows: s (singlet), d (doublet), t (triplet), q (quartet), dd (double doublet), p (pentet), sp (septet), m (multiplet), br. (broad).

The mixture of equimolar amounts of amino acid and sodium cyanate (NaOCN) in water was kept at a room temperature for 75-80 hours (Compounds: **2, 4, 7-13**) or at 100 °C for 4 hours (Compounds: **3, 5, 6**). Then pH of reaction mixture was adjusted to 2 – 3 with concentrated HCl. The separated solid was filtered, washed with water, and recrystallized. From filtrate additional amount of product was obtained after concentrating at reduced pressure. Reaction **14** was carried out in 75% ethanol (100 °C, 4 h). After removing ethanol under reduced pressure, water was added, and pH was adjusted to 5-6 with concentrated HCl. The separated product was treated as above.

### RrCβAA Activity assay

RrCβAA assays were performed at 40°C. The reaction mixture contained 100 mM phosphate buffer (pH 8.0), and 5 μg purified enzyme in a total volume of 0.1 mL. Reactions were started by the addition of 0.1 mL N-carbamoyl-L-β-alanine to final concentrations of 100 mM after preincubation of both reaction mixture and substrate solutions at 40 °C for 10 min. After 15 minutes the reaction was stopped by adding 30 % w/v trichloroacetic acid (TCA) to a final concentration of 3 % w/v. Specific activity of RrCβAA was determined using an assay able to detect β-alanine concentration, upon conversion into an isoindole derivative by reaction with ortho-phthalaldehyde (OPA) [44]. Particular attention was paid to the OPA reaction conditions, as it has been reported that the derivative of β-alanine is unstable. For this reason, a high concentration of reagents (20-times excess of OPA and 50-times excess of β-mercaptoethanol compared to the β-alanine product) was used to stabilize the final product. Thus, 3 mL of freshly prepared activity reagent (0.1 M sodium borate pH 9.6, 2.5 mM OPA and 2.5 mM β-mercaptoethanol) were added to each sample, followed by incubation at 20 °C for 30 min. β-alanine concentrations were determined spectrophotometrically at SF-46 (“Lomo”, Russia) based on the absorption of the corresponding isoindole at 340 nm. The extinction coefficient for each substrate was calculated separately (extinction coefficient data is avaible in **Supplementary Figure 7**. For preparation of standard curves 40 mM concentration of L-β-alanine, L-α-alanine, L-α-valine, L-β-phenyl-α-alanine, α-amino butyric acid, γ-amino butyric acid, L-α-leucine, L-α-methionine and glycine was prepared. And the adsorption of serial dilutions of amino acids at final concentrations of 0.032 mM, 0.064 mM, 0,096 mM, 0.128 mM, and 0.16 mM, were measured. One unit of enzyme activity was defined as the amount of enzyme catalysing the formation of one micromole of product per minute under the above mentioned conditions. Specific activity was calculated per milligram of protein. All measurements were done at least in two separate experiments with two replicates.

### RrCβAA temperature optimum and thermostability

For determination of the optimal temperature for RrCβAA, enzymatic activity was measured under the described conditions at various temperatures ranging from 25 to 70°C. Thermostability of purified RrCβAA was investigated by incubating RrCβAA at various temperatures (25–70°C) for 15 min in phosphate buffer, followed by incubation on ice. Residual activities were determined under the above assay conditions.

### Effect of metals and chelation on RrCβAA activity

Metal ions are generally considered as important factors affecting microbial enzyme activity. The effects of various mono- and bivalent metal ions (including NaCl, KCl, CaCl_2_, MgSO_4_, BaCl_2_, SnCl_2_, PbSO_4_, FeCl_3_, FeSO_4_, CuSO_4_, ZnSO_4_, MnSO_4_, CdCl_2_, NiCl_2_, CoSO_4_,) and chemical compounds (including EDTA, 5,5′-dithiobis-(2-nitrobenzoic acid) (DTNB), dithiothreitol (DTT), β-mercaptoethanol) on RrCβAA activity was investigated. RrCβAA was incubated in the presence of 2 mM of each metal ion, DTT and EDTA, or 5 mM of DTNB and β-mercaptoethanol, for one hour at 4°C. A control was performed in the absence of any tested compound. To test recovery of enzyme activity after metal removal, the enzyme was incubated with 5 mM EDTA at 4°C for one hour to chelate metals, then dialysed against excess reaction buffer containing 2 mM of each metal ion tested. All the activity assays were performed in triplicate.

### Substrate spectrum and enantioselectivity of RrCβAA

The specific activity of purified RrCβAA toward various N-carbamoyl-amino acids including N-carbamoyl-L-β-alanine, N-carbamoyl-L-α-alanine, N-carbamoyl-D-α-alanine, N-carbamoyl-DL-α-alanine, N-carbamoyl-L-α-valine, N-carbamoyl-D-α-valine, N-carbamoyl-DL-α-valine, N-carbamoyl-L-β-phenyl-α-alanine, N-carbamoyl-L-β-phenyl-β-alanine, N-carbamoyl-D-β-phenyl-α-alanine, N-carbamoyl-DL-β-phenyl-α-alanine, N-carbamoyl-α-amino butyric acid, N-carbamoyl-γ-amino butyric acid, N-carbamoyl-L-α-leucine, N-carbamoyl-L-α-methionine, and N-carbamoyl-α-glycine was measured using the above method. A calibration curve for each product was constructed and extinction coefficient for each product has been calculated (**Supplementary Figure 7**). Neither the isoindole formed from ammonium ions, nor N-carbamoyl-amino acids gave a detectable signal under the chosen reaction conditions.

### Protein quantification

The concentration of the purified was determined by a colorimetric technique using the PierceTM BCA protein assay kit following manufacturer’s specifications for the standard test-tube procedure at 37°C. Diluted bovine serum albumin (BSA) standards were prepared in GF buffer and a calibration curve of absorbance at 562 nm against concentration was plotted (**Supplementary Figure 8**). Protein sample absorbance was measured at 562 nm (average of three experimental replicates) and the concentration was calculated.

### Protein Crystallography

Purified recombinant RrCβAA was concentrated to 15 mg/mL using a 10 kDa MWCO centrifugal concentrator (Vivaspin) and subjected to sitting drop vapor diffusion crystallization screening with commercial screens from Molecular Dimensions and Hampton Research. Drops of 100 nL protein plus 100 nL well solution were set up against wells containing 70 μL of crystallisation solutions. After two weeks crystals were found in row D of the PACT premier screen (Molecular Dimensions). An optimisation screen based on this condition was set up in 24 well plates by varying the PEG1500 concentration and MMT buffer pH. Drops of 1 μL protein and 1 μL well solution were set up on plastic cover slips over wells containing 1 mL crystallisation solution. Crystals grew in a well solution containing 23 % (w/v) PEG1500 and 100 mM MMT pH 6.0. Crystals were harvested with a LithoLoop (Molecular Dimensions Limited) and transferred to a cryoprotection solution of well solution supplemented with 50 % (v/v) PEG400. Cryoprotected crystals were flash cooled in liquid nitrogen. Diffraction data were collected at Diamond Light Source; data collection and model refinement statistics are shown in **Supplementary Table 7**. Diffraction data are available at <u>doi:10.5281/zenodo.7331274</u>.

The data set was integrated with XIA2 [45] using DIALS [46],and scaled with Aimless [47]. The space group was confirmed with Pointless [48]. The phase problem was solved with MorDa. Initial model building was performed with CCP4build task on CCPcloud [49]. The model was refined with iterative cycles of refmac[50], or BUSTER, intercalated with manual model building with COOT [51]. The model was validated using Coot and Molprobity [52]. Other software used were from CCP4 cloud and the CCP4 suite [53]. Structural figures were produced with ChimeraX [54].

## Acknowledgements

The authors would like to thank Diamond Light Source for beamtime (proposal mx18598), and the staff of beamline I04-1.

## Author Contributions

Study conceptualization: AP, MK, SK, GA

Investigation: Initial microbial strain identification and characterisation: MK; Molecular Biology – AP; Protein purification and characterisation – AP, AS, MK, CP, WAS; Structural Biology – AB, JMW; Preparation and validation of Substrates: – MDK, KD, HP

Resources – JMW, GA, AP, AB

Funding acquisition – GA, AP, JMW

Writing – Original Draft: AP, JMW

Writing – Review and Editing: AP, AH, JMW, AB, AK, GA

Visualisation – AP, AS, JMW, AB

## Data availability Statement

All data used to prepare this manuscript are available as supplementary materials or deposited at publicly accessible databases. Links and references to datasets are in the Methods and Supplementary Materials.

