## Supplementary Information for "Structural and biochemical characterisation of the N-Carbamoyl-β-Alanine Amidohydrolase from *Rhizobium radiobacter* MDC 8606"

* To whom correspondence should be addressed

### Contents

**Supplementary Figures**

Supplementary Figure 1. Annotated sequence alignment of carbamoylase enzymes

Supplementary Figure 2. Immobilised metal affinity purification of RrCβAA

Supplementary Figure 3. RrCβAA dimerization interface

Supplementary Figure 4. Rotation of the RrCβAA catalytic domain at the dimerization domain boundary

Supplementary Figure 5. Presence of electron density consistent with a MES buffer molecule in the RrCβAA active site

Supplementary Figure 6. NMR spectra of ligands produced for this study

Supplementary Figure 7. OPA derivatised amino acid standard curves for quantification of millimolar extinction coefficients

Supplementary Figure 8. Bovine serum albumin standard curves for protein quantification by BCA.

**Supplementary Tables**

Supplementary Table 1. Purification table of recombinant RrCβAA

Supplementary Table 2. Temperature - activity relationship for RrCβAA

Supplementary Table 3. Temperature - stability relationship for RrCβAA

Supplementary Table 4. RrCβAA activity in the presence of divalent cations and reducing agents

Supplementary Table 5. Activity recovery of EDTA inactivated RrCβAA enzyme

Supplementary Table 6. Substrate preference of RrCβAA enzyme

Supplementary Table 7. X-ray crystallography data collection and refinement statistics

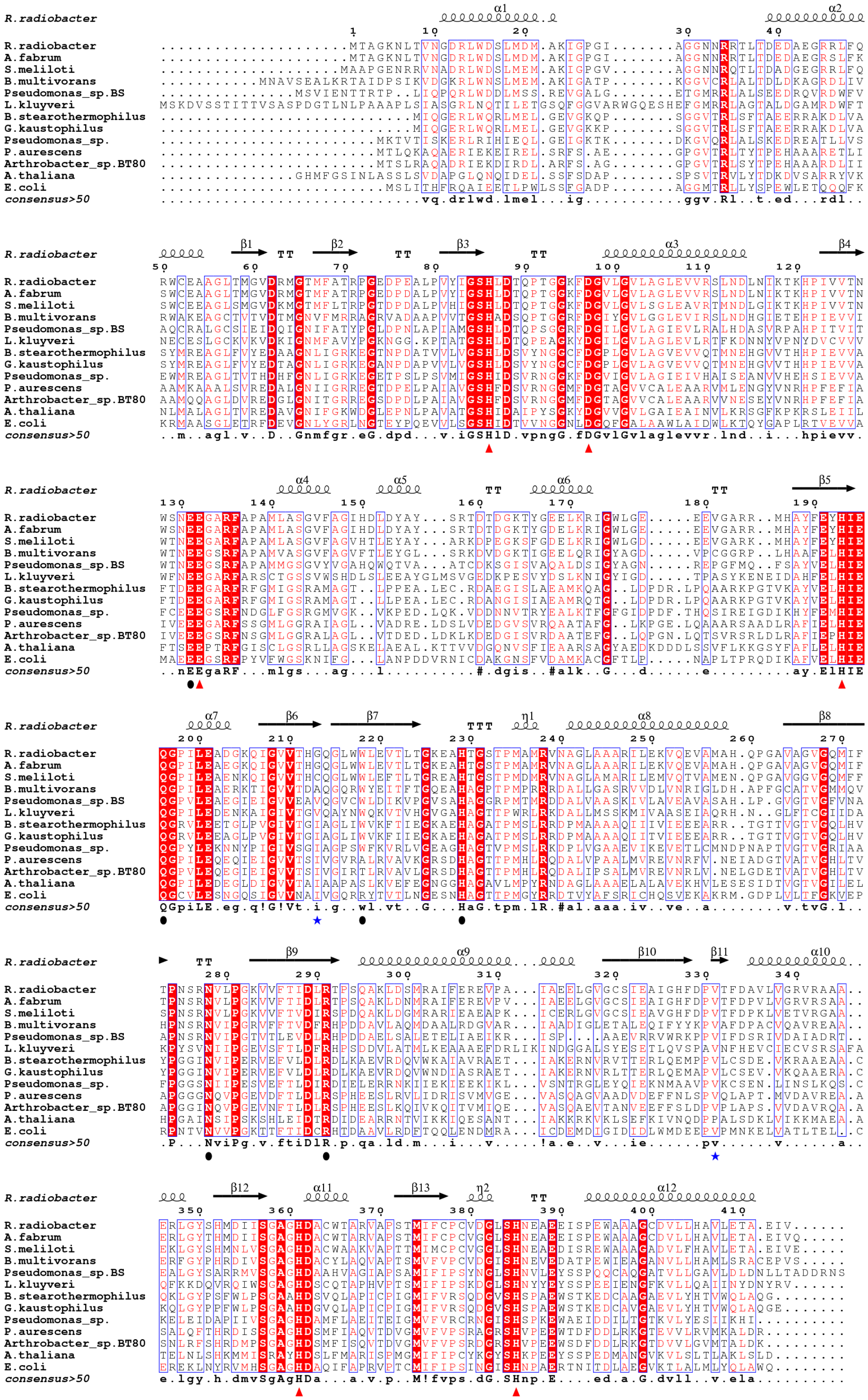

**Supplementary Figure 1. Annotated sequence alignment of carbamoylase enzymes.** A multiple sequence alignment of carbamoylase enzymes was performed using Multalin and visualised with ESPript. Secondary structure elements from the crystal structure of the R. radiobacter enzyme are shown above the alignment. Strictly conserved residues are shown with a red background, while partially conserved residues are shown with red text. Conserved residues involved in metal coordination are highlighted with red triangles, residues involved in ligand binding with black ovals and hinge residues between the active site and dimerisation domains are shown with blue stars.

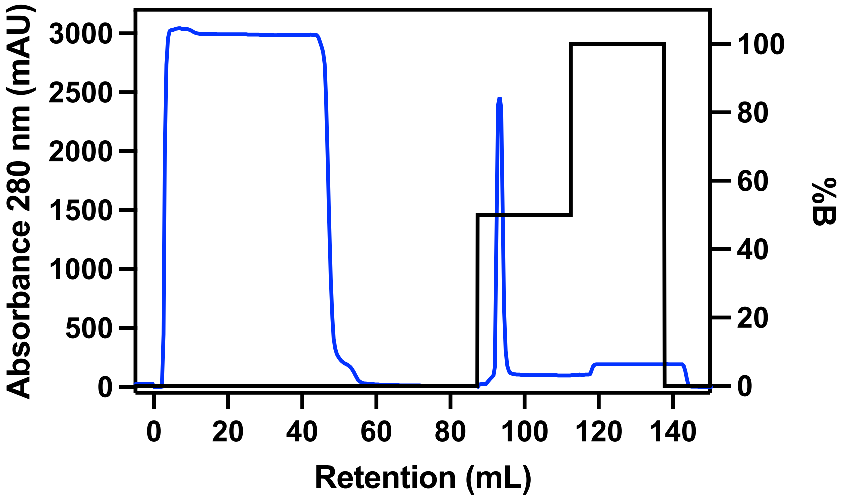


**Supplementary Figure 2. Immobilised metal affinity purification of RrCβAA.** A) UV Chromatogram of RrCβAA purification by immobilised metal affinity chromatography on an Åkta Pure chromatography system. The sample was loaded onto a column equilibrated with buffer HisA (50 mM Tris.HCl, pH 8, 500 mM NaCl, 50 mM Imidazole) and eluted with a step gradient of buffer HisB (50 mM Tris.HCl, pH 8, 500 mM NaCl, 500 mM Imidazole). The sharp peak at 95 mL corresponds to the RrCβAA protein.

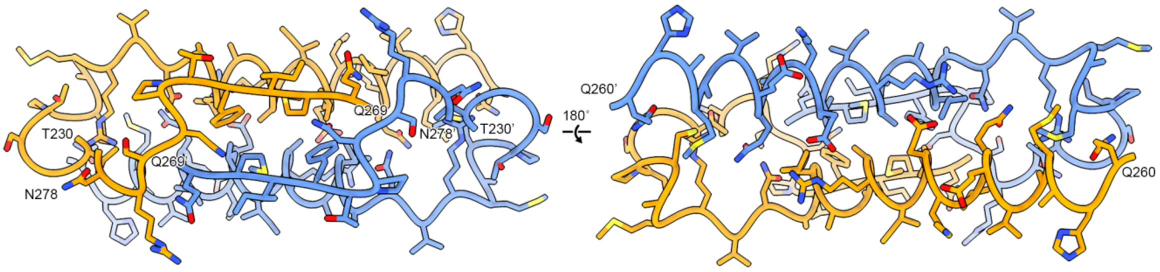


**Supplementary Figure 3. RrCβAA dimerization interface.** The RrCβAA dimerization interface is shown as cartoon backbone representation with interacting residues shown as sticks coloured by atom. The subunits are coloured orange and blue, with residues from the blue subunit indicated with a prime symbol.

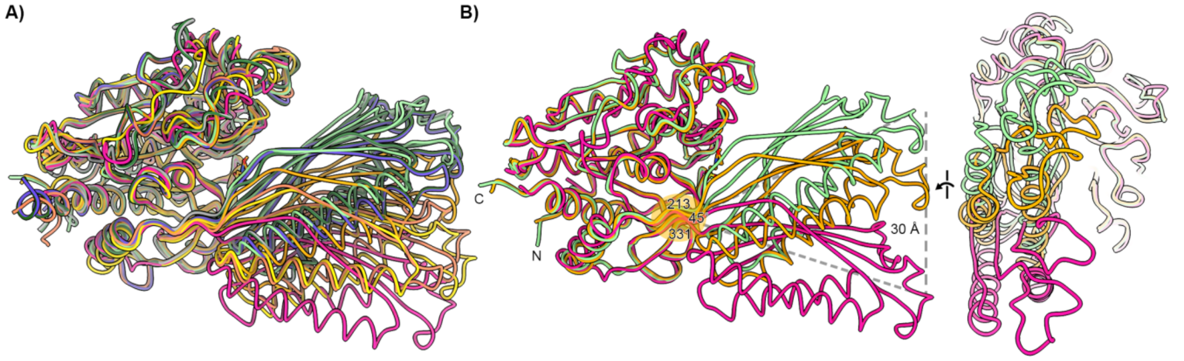


**Supplementary Figure 4. Rotation of the RrCβAA catalytic domain at the dimerization domain boundary.** A) Range of rotation present in published homologue structures, shown as licorice representation. Models are coloured as follows: 8APZ – mint; 5THW – peach; 3N5F – pink;1R3N – salmon; 1Z2L – yellow; 2V8G – purple; 4PXB – grey; RrCβAA – Orange. B) Extremes of rotation between the domains highlighted with 8APZ and 3N5F – the hinge residues in RrCβAA are highlighted with the maximum angle of rotation and extent of relative displacement of the catalytic domain to the dimerization domain.

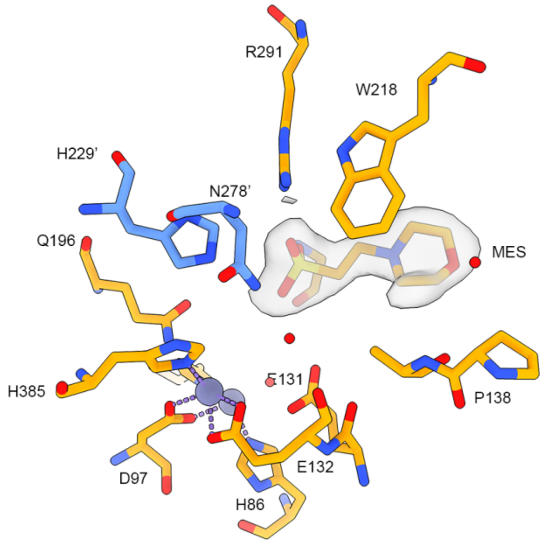


**Supplementary Figure 5. Presence of electron density consistent with a MES buffer molecule in the RrCβAA active site.** Based on features in the 2mFo-DFc electron density map a MES buffer molecule was modelled in the active site. Residues interacting with the MES molecule are shown as stick representation in orange and blue, with the carbon atoms of the MES molecule shown in orange. Final 2mFo-DFc Electron density map rendered at 1σ as a transparent grey surface.

**Supplementary Figure 6. NMR spectra of ligands produced for this study.**

**Compound 1: 4-(Methylthio)-2-ureidobutanoic acid.** From L-methionine. M. p. 182-183 °C (EtOH-$H_2O$, 5:1). $^1H$ NMR δ: 1.74 – 1.86 (m, 1H) and 1.92 – 2.03 (m, 1H, both $CH_2$); 2.06 (s, 3H, $CH_3$); 2.47 (t, 2H, J = 7.7 Hz, $SCH_2$); 4.19 (td, 1H, J = 8.2, 4.9 Hz, CH); 5.50 (br., 2H, $NH_2$); 6.30 (d, 1H, J = 8.2 Hz, NH); 12.52 (br., 1H, COOH). $^{13}C$ NMR δ: 14.7 ($CH_3$); 29.6 ($CH_2$); 31.9 ($CH_2$); 51,4 (CH); 158.4 (NCO); 174.1 (OCO).

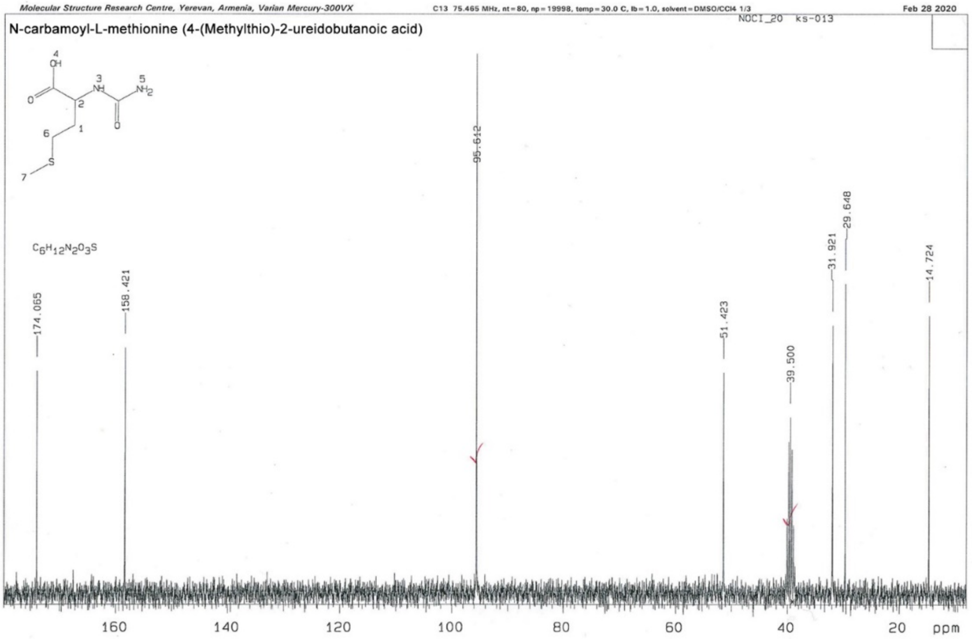


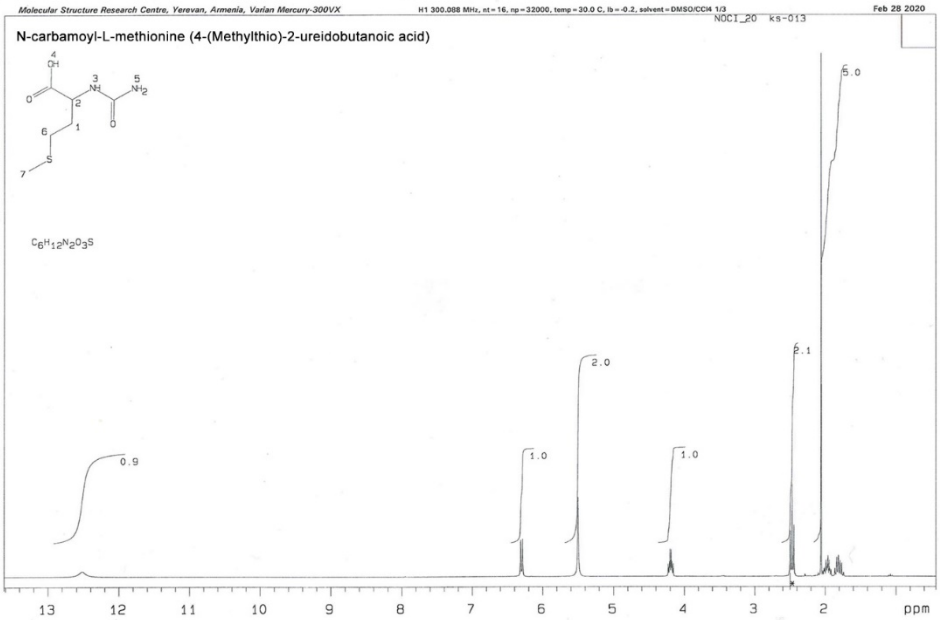

**Compound 2: 4-Methyl-2-ureidopentanoic acid.** From L-leucine.M. p. 227-230 ºC ($H_2O$) (lit.: 216-218 ºC) [1]. $^1$H NMR δ: 0.92 (d, 3H, J = 6.5 Hz) and 0.93 (d, 3H, J = 6.5 Hz, both 2$CH_3$); 1.37 – 1.55 (m, 2H, $CH_2$); 1.63 – 1.80 (m, 1H, $CH_3C$**H**$CH_3$); 4.11 (ddd, 1H, J = 8.9, 8.6, 5.3 Hz, CH); 5.40 (br., 2H, $NH_2$); 6.12 (d, 1H, J = 8.6 Hz, NH); 12.25 (v.br., 1H, COOH). $^{13}$C NMR δ: 21.6 ($CH_3$); 22.7 ($CH_3$); 24.2 (CH); 41.3 ($CH_2$); 50.6 (NCH); 158.3 (NCO); 175,1 (OCO).

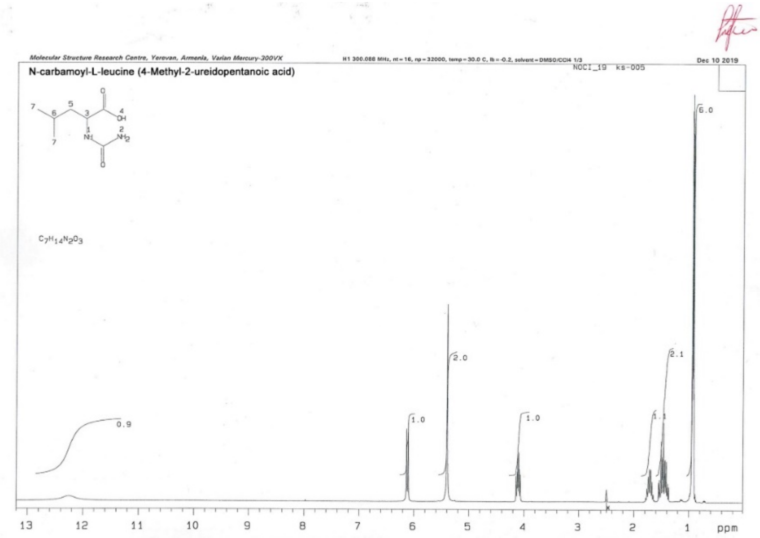


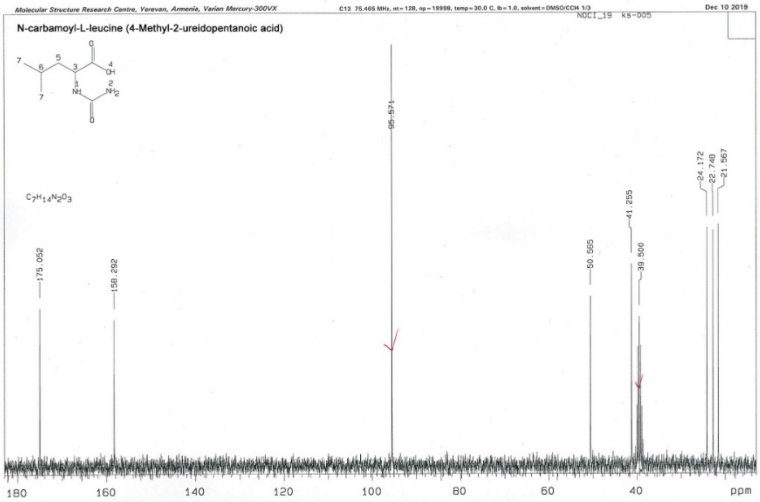


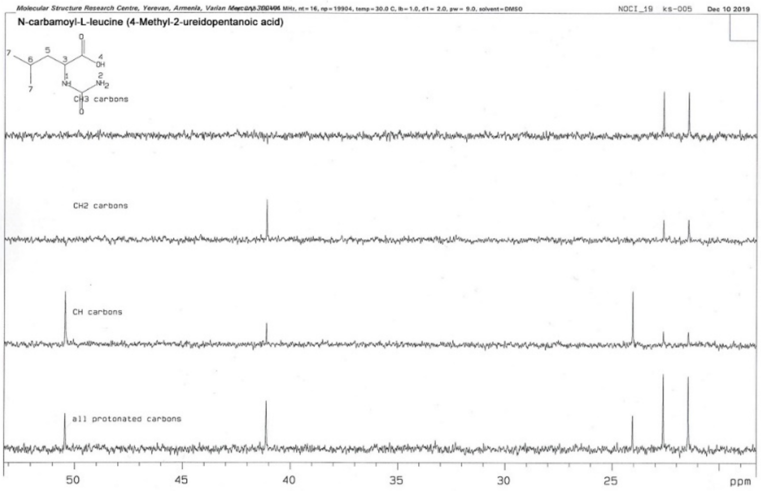

**Compound 3: 4-Ureidobutanoic acid**. From 4-aminobutanoic acid. M. p. 187-188 °C (EtOH-$H_2O$, 5:1), lit.: 174-175 °C [ 2 ], 178-179 °C [ 3 ]. $^1$H NMR δ: 1.65 (tt, 2H, J = 7.4, 6.7 Hz, $CH_2$); 2.21 (t, 2H, J = 7.4 Hz, O=C-$CH_2$); 3.02 (td, 2H, J = 6.7, 5.8 Hz, $NCH_2$); 5.21 (br., 2H, $NH_2$); 5.91 (br. t, 1H, J = 5.8 Hz, NH); 11.88 (br., 1H, COOH). $^{13}$C NMR δ: 25.4 ($CH_2$); 30,9 ($CH_2$); 38.3 ($CH_2$); 158.6 (NCO); 173.9 (OCO).

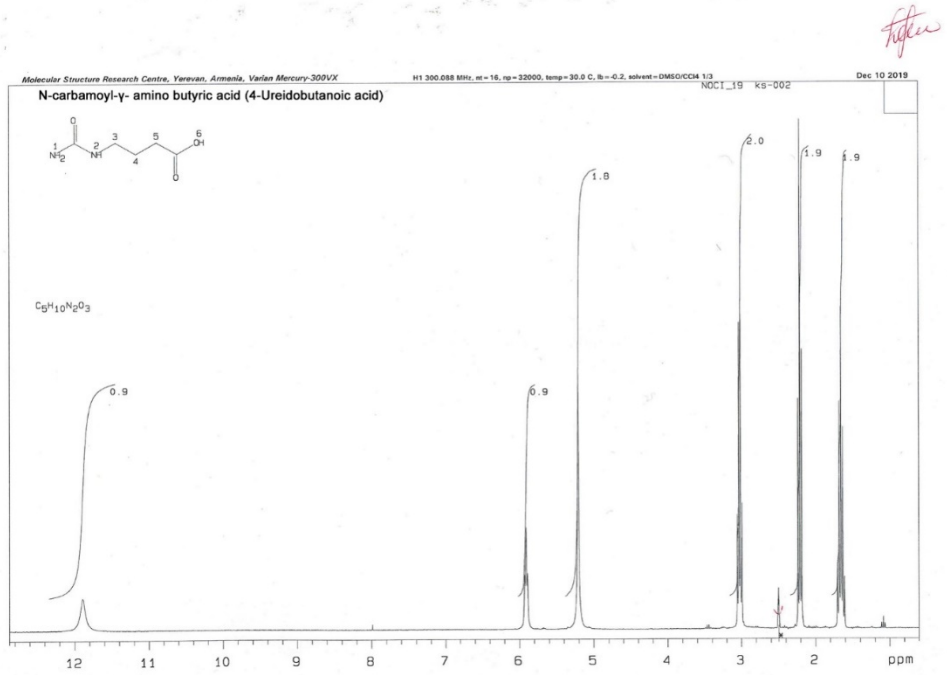


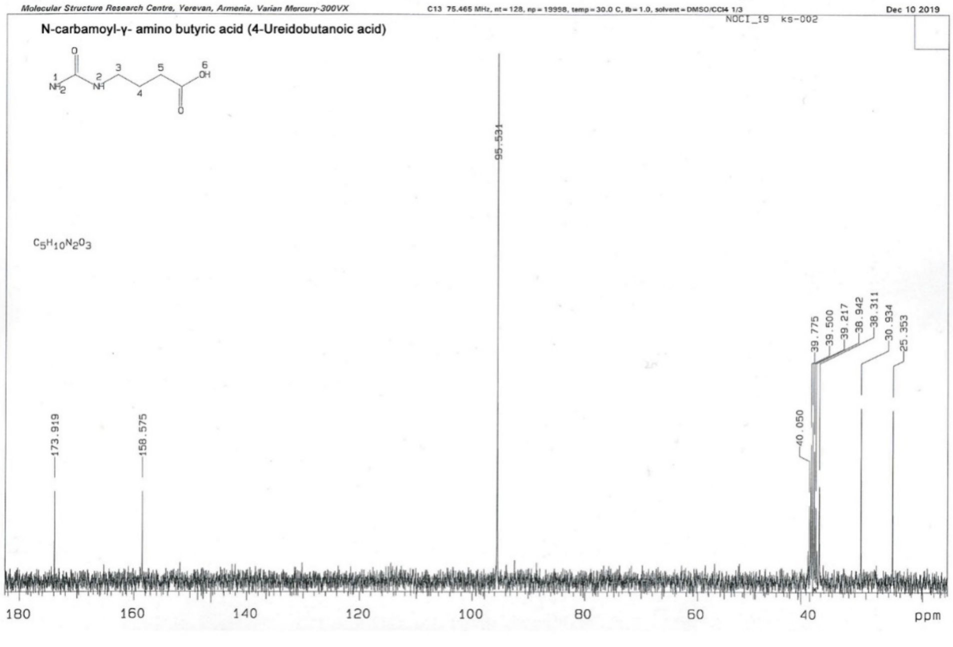

**Compound 4: 2-Ureidobutanoic acid.** From DL-2-aminobutanoic acid. M. p. 196-197 °C ($EtOH-H_2O$, 5:1), lit.: 178-179 °C [ 3 ]. $^{1}$HNMR δ: 0.91 (t, 3H, J = 7.4 Hz, $CH_3$); 1.53 – 1.80 (m, 2H, $CH_2$); 4.06 (ddd, 1H, J = 7.83, 7.1, 5.3 Hz, CH); 5.50 (br., 2H, $NH_2$); 6.22 (d, 1H, J = 8.3 Hz, NH); 12.40 (br., 1H, COOH). $^{13}$C NMR δ: 9.6 ($CH_3$); 25.3 ($CH_2$); 53.3 (CH); 158.5 (NCO); 174.4 (OCO).

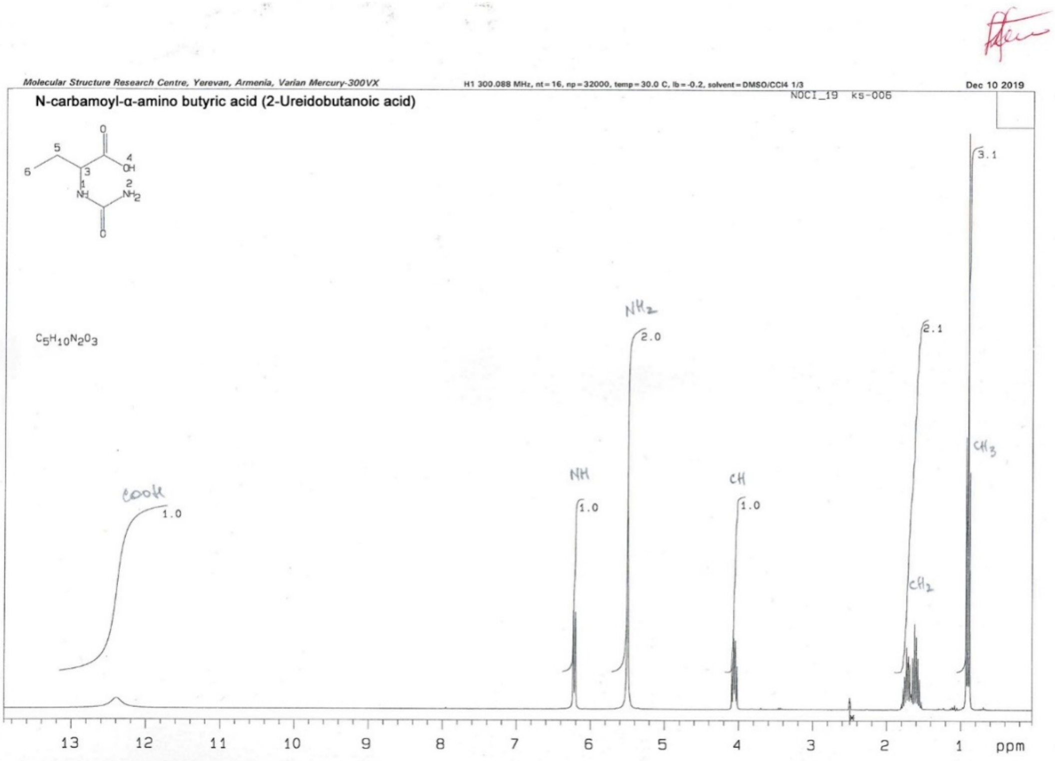


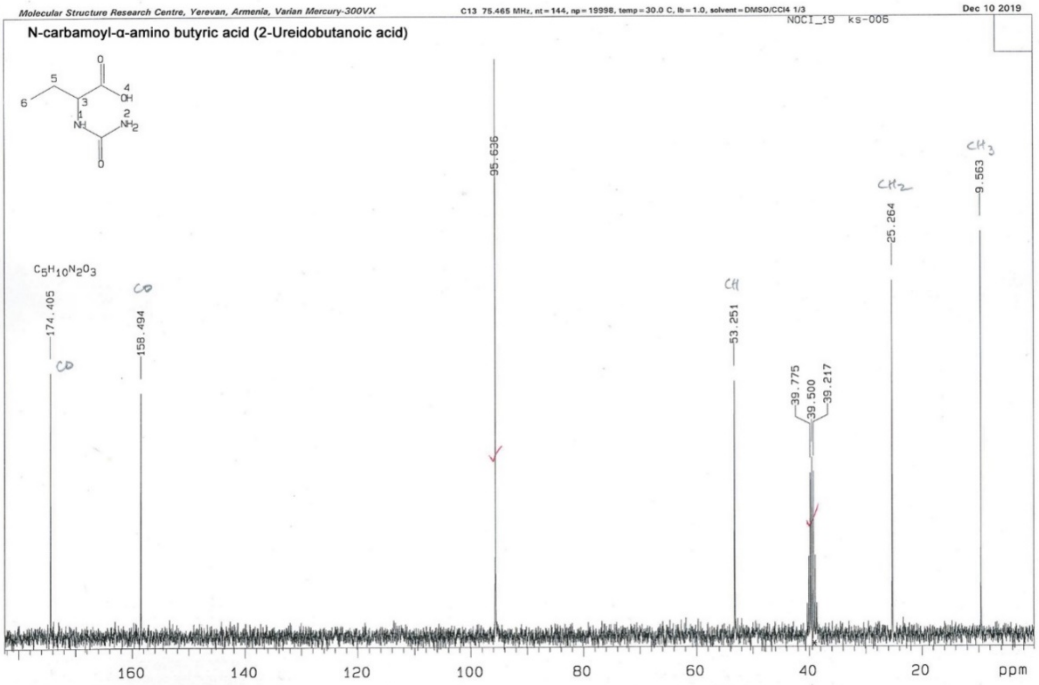

**Compound 5: 3-Phenyl-2-ureidopropionic acid.** From DL-phenylalanine. M. p. 198 °C ($H_2O$). $^1$H NMR δ: 2.93 (dd, 1H, J = 13.7, 7.0 Hz) and 3.04 (dd, 1H, J = 13.7, 5.4 Hz, both $CH_2$); 4.41 (ddd, 1H, J = 8.2, 7.0, 5.4 Hz, CH); 5.46 (br., 2H, $NH_2$); 6.14 (d, 1H, J = 8.2 Hz, NH); 7.13 – 7.28 (m, 5H, Ar); 12.41 (v.br., 1H, COOH).$^{13}$C NMR δ: 37.2 ($CH_2$); 52.9 (CH); 125.3 (CH); 127.1 (2CH); 128.6 (2CH); 136.7; 157.4 (NCO); 173.0 (OCO).

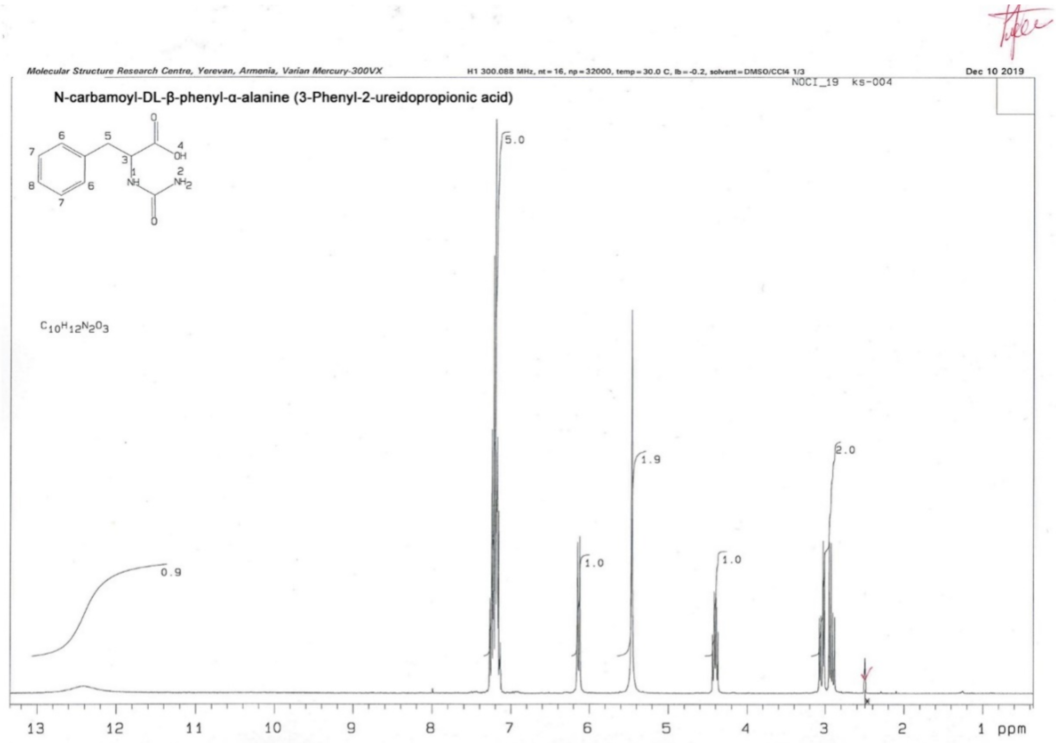


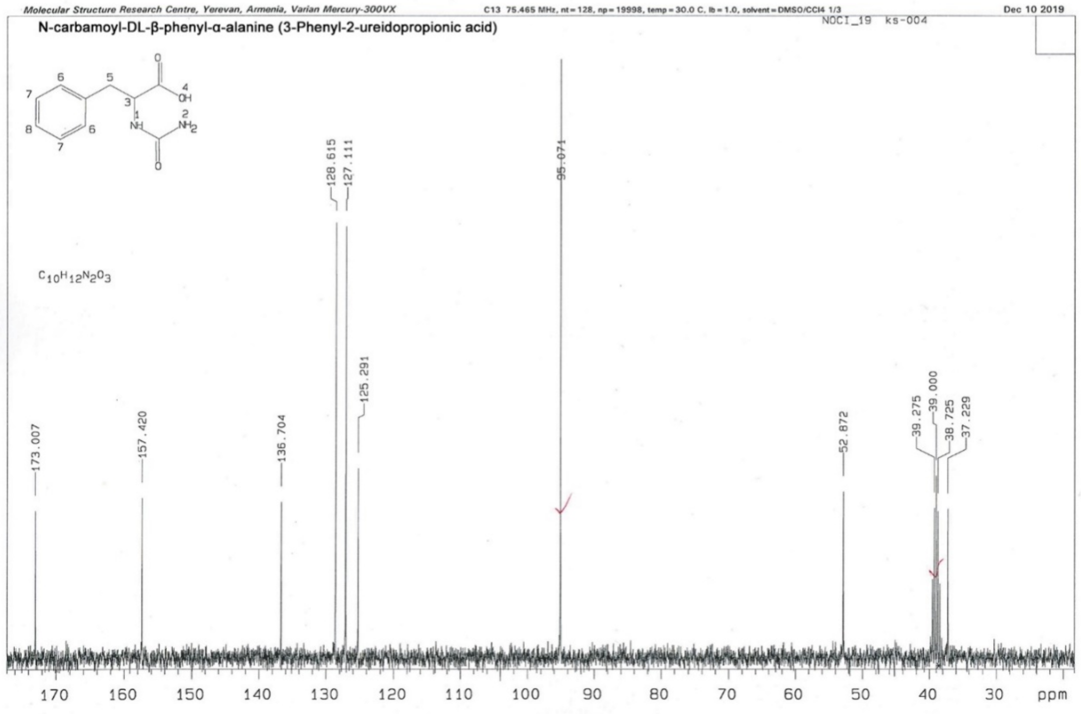

**Compound 6: 3-Phenyl-2-ureidopropanoic acid.** From L-phenylalanine. M. p. 210-211 °C ($H_2O$) (lit.: 200 °C) [1]. Data of $^1H$ and $^{13}C$ NMR are same as for **5**.

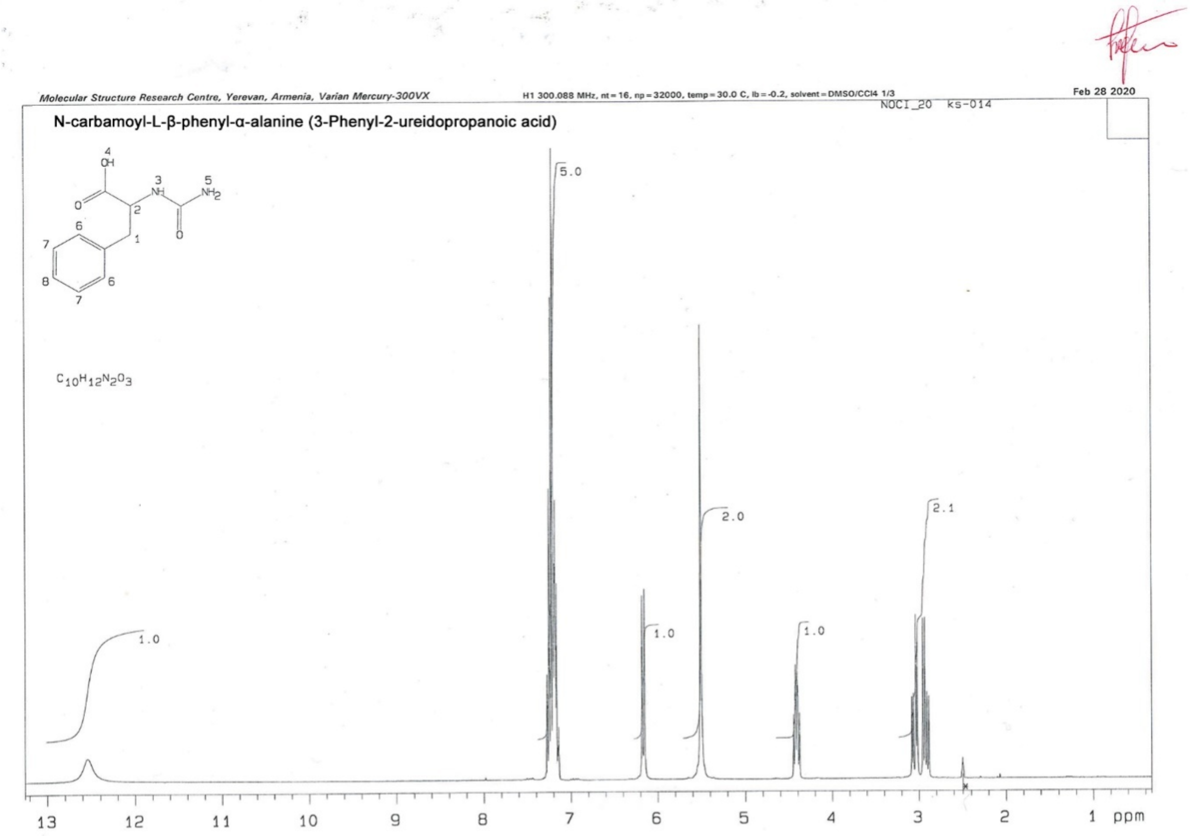


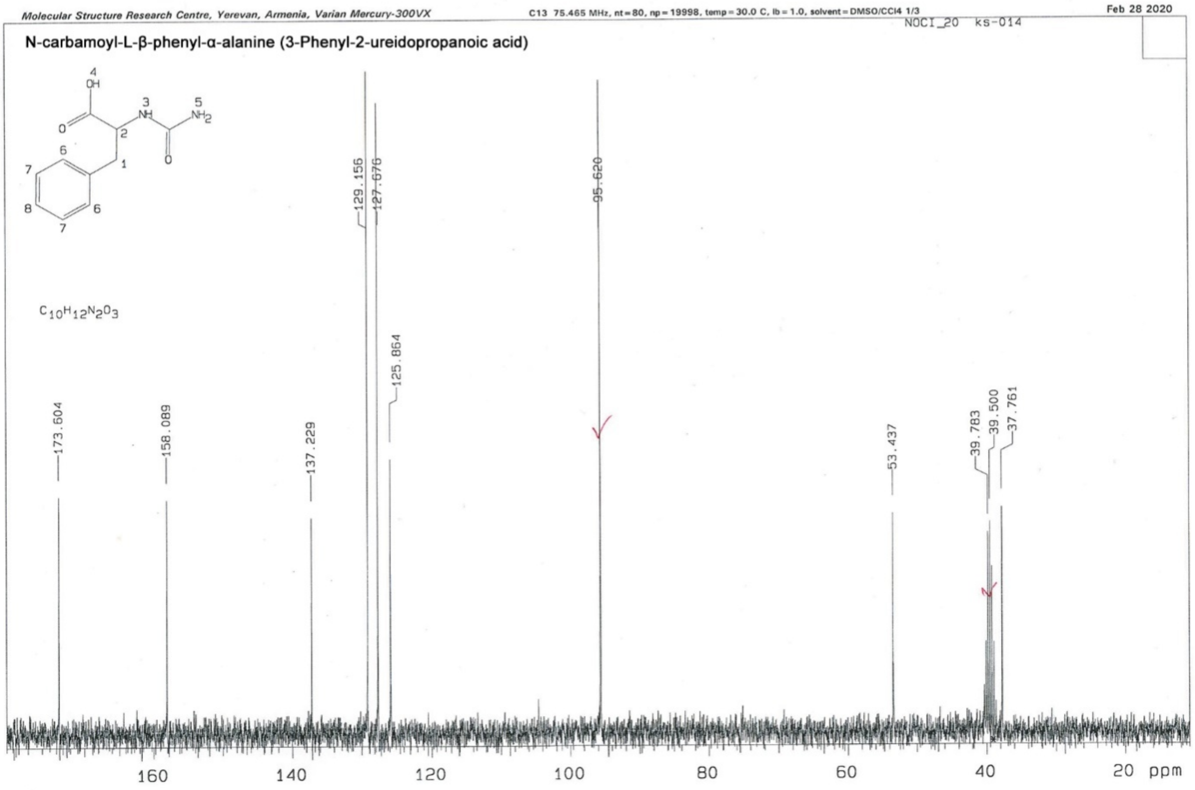

**Compound 7: 3-Phenyl-3-ureidopropanoic acid.** From DL-3-amino-3phenylpropanoic acid. M. p. 203-206 °C (EtOH). $^1$H NMR δ: 2.62 (dd, 1H, J = 15.3, 6.3 Hz) and 2.66 (dd, 1H, J = 15.3, 7.6 Hz, both $CH_2$); 5.05 (ddd, 1H, J = 8.6, 7.6, 6.3 Hz, CH); 5.39 (br. 2H, $NH_2$); 6.55 (d, 1H, J = 8.6 Hz, NH); 7.15-7.33 (m, 5H, Ar); 12.10 (br. 1H, COOH). $^{13}$C NMR δ: 41.4 ($CH_2$); 49.9 (CH); 126.0 (2CH); 126.2 (CH); 127.7 (2CH); 143.2; 157.8 (NCO); 171.8 (OCO).

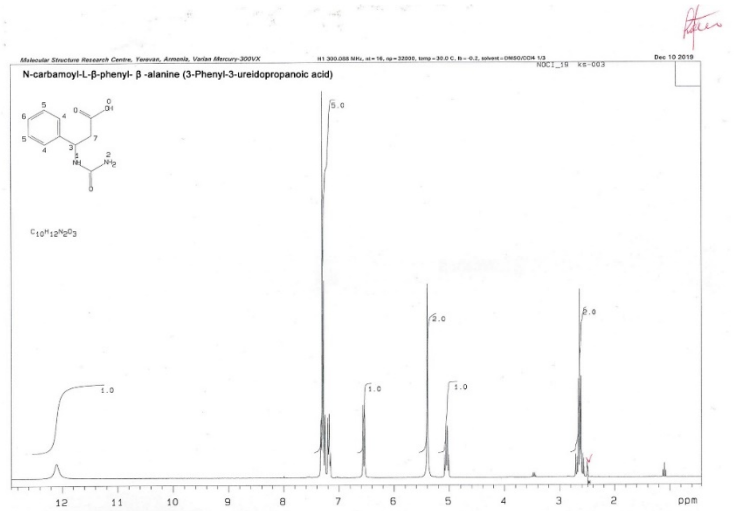


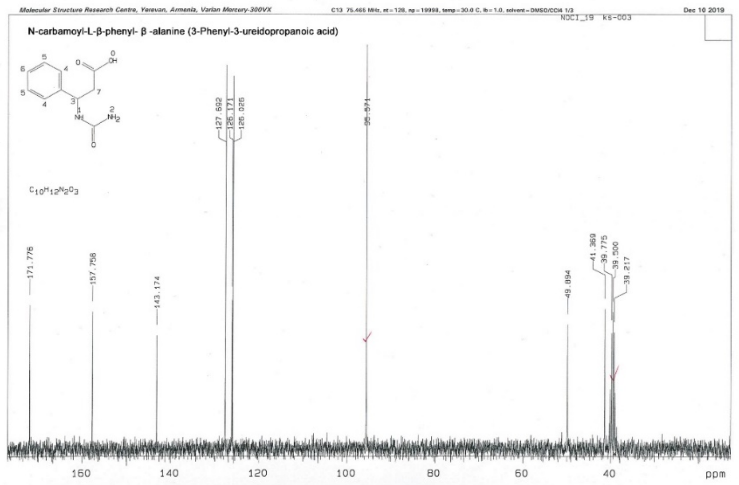


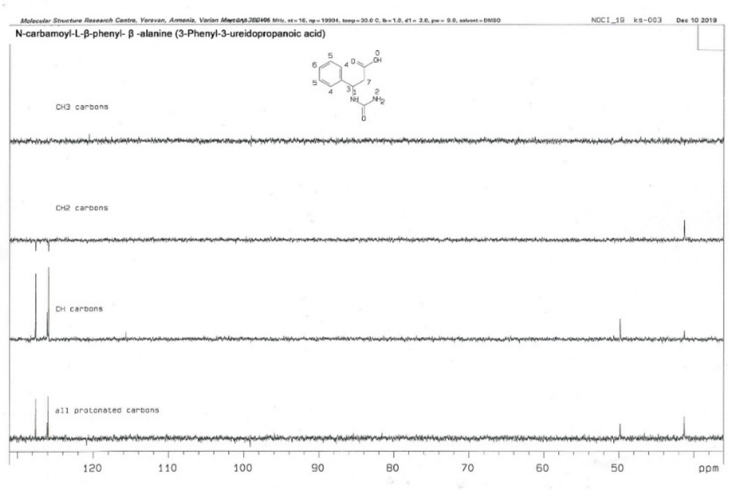

**Compound 8: 3-Methyl-2-ureidobutanoic acid.** From DL-valine. M. p. 211-213 °C (EtOH-$H_2O$, 5:1). M. p. 218-220 °C (EtOH-$H_2O$, 5:1), lit.: 207-209 °C [1]. $^1$H NMR δ: 0.88 (d, 3H, J = 6.8 Hz) and 0.93 (d, 3H, J = 6.8 Hz, both $2CH_3$); 2.05 (dsp, 1H, J = 6.8, 4.9 Hz, $CH_3C\mathbf{H}CH_3$); 4.05 (dd, 1H, J = 9.1, 4.9 Hz, NCH); 5.43 (br., 2H, $NH_2$); 6.13 (br. d, 1H, J = 9.1 Hz, NH); 12.28 (br., 1H, COOH). $^{13}$C NMR δ: 17.4 ($CH_3$); 18.9 ($CH_3$); 30.3 (CH); 57.0 (NCH); 158.4 (NCO); 173.9 (OCO).

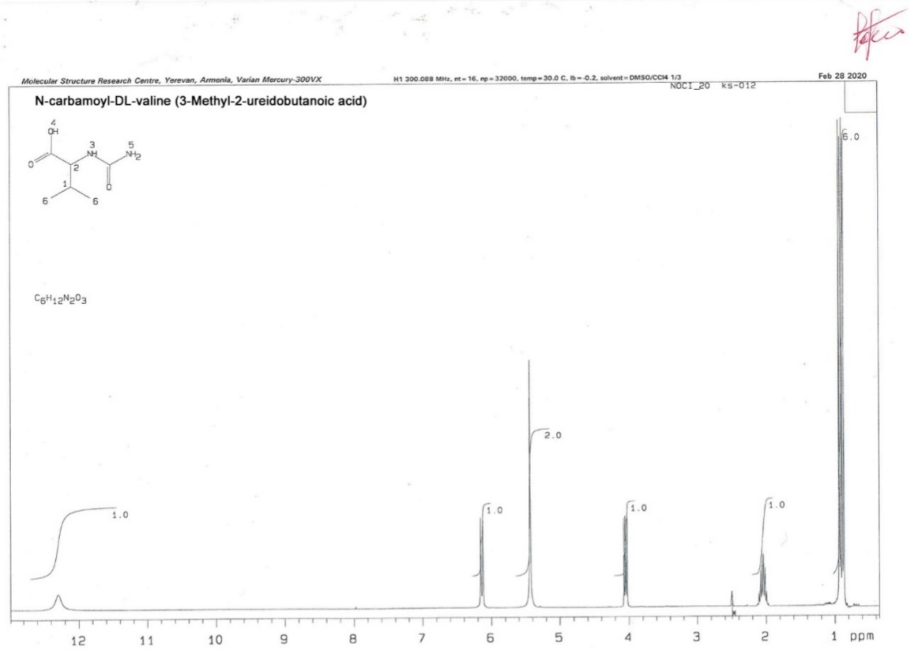


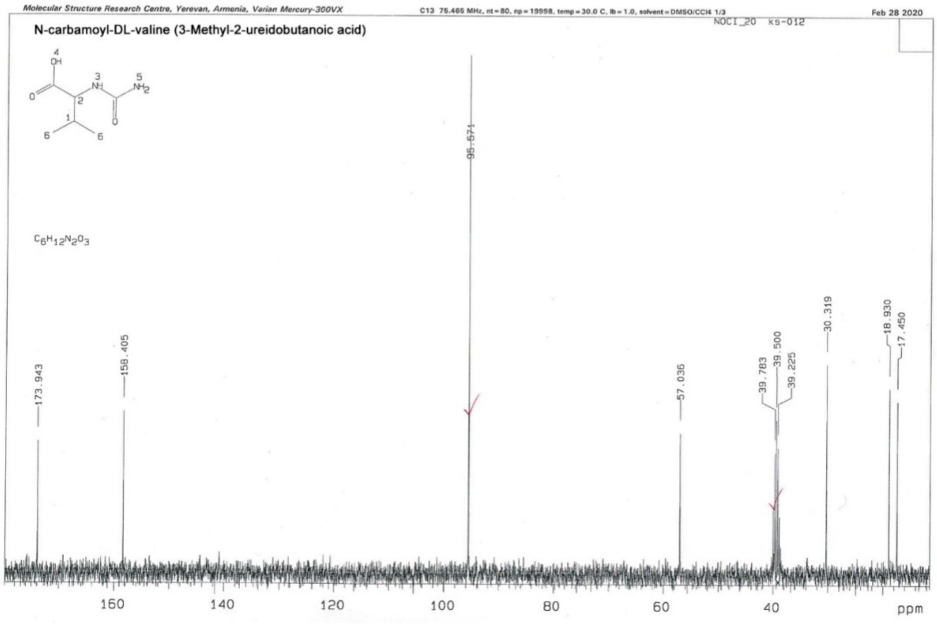

**Compound 9: 3-Methyl-2-ureidobutanoic acid.** From D-valine. M. p. 225-226 °C (EtOH-$H_2O$, 5:1). Data of $^1H$ and $^{13}C$ NMR are same as for **8**.

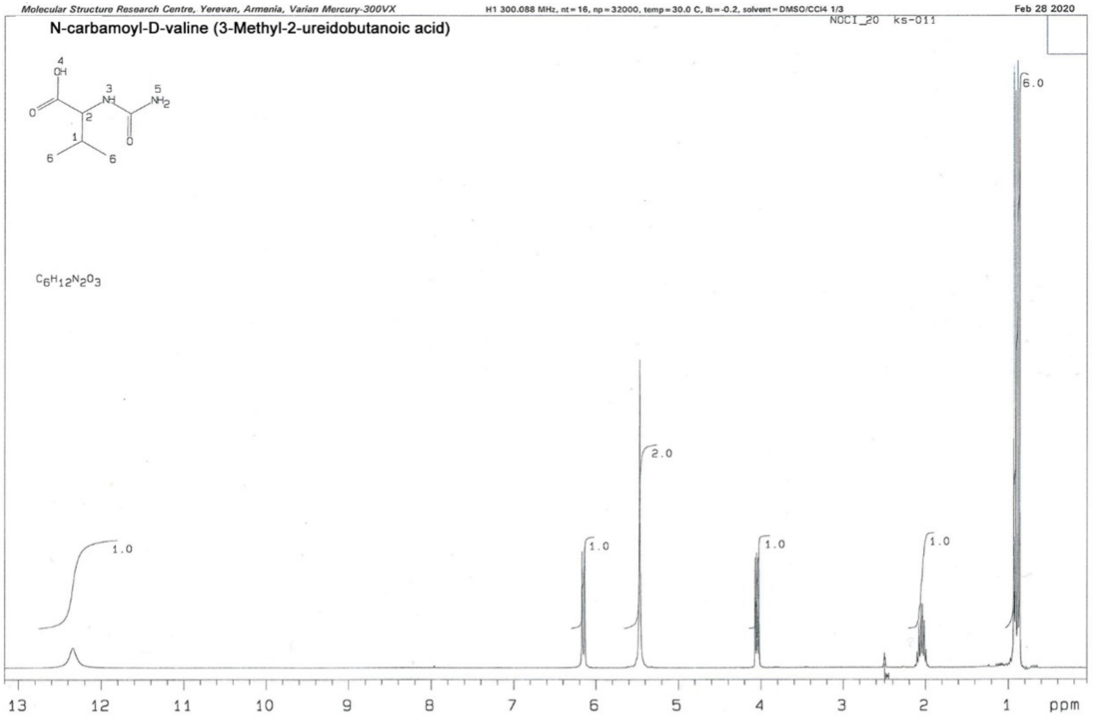


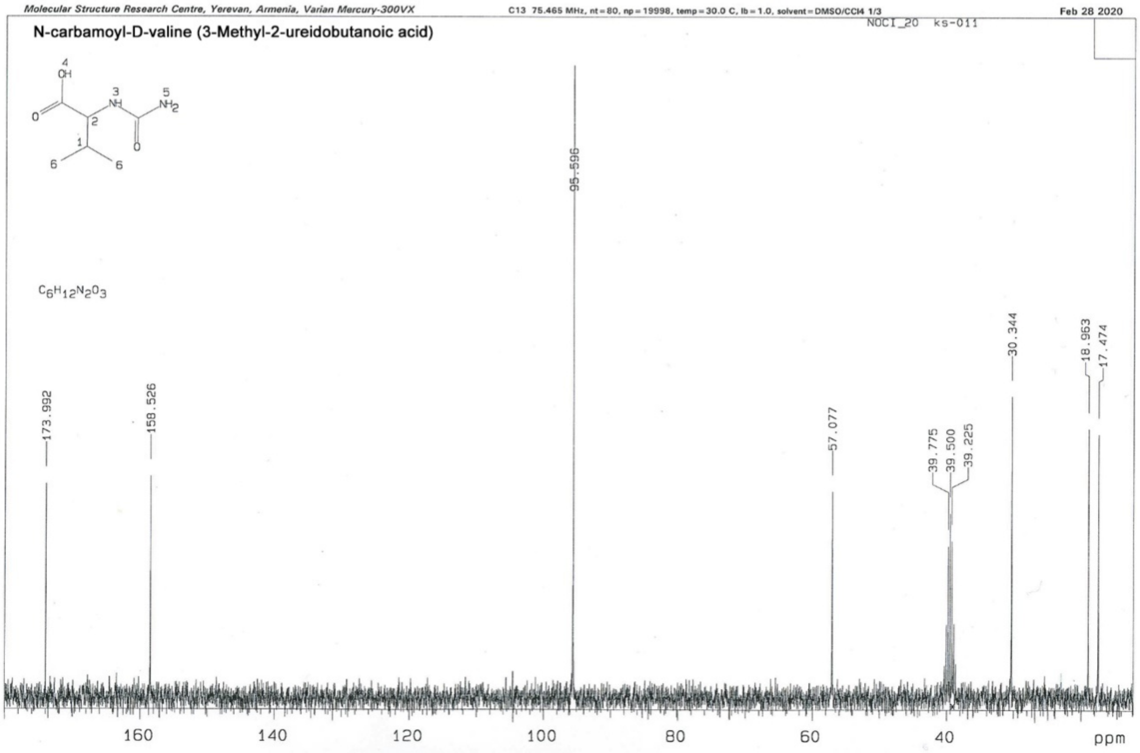

**Compound 10: 3-Methyl-2-ureidobutanoic acid.** From L-valine. M. p. 218-220 °C (EtOH-$H_2O$, 5:1), lit.: 207-209 °C [1]. Data of $^1H$ and $^{13}C$ NMR are same as for **8.**

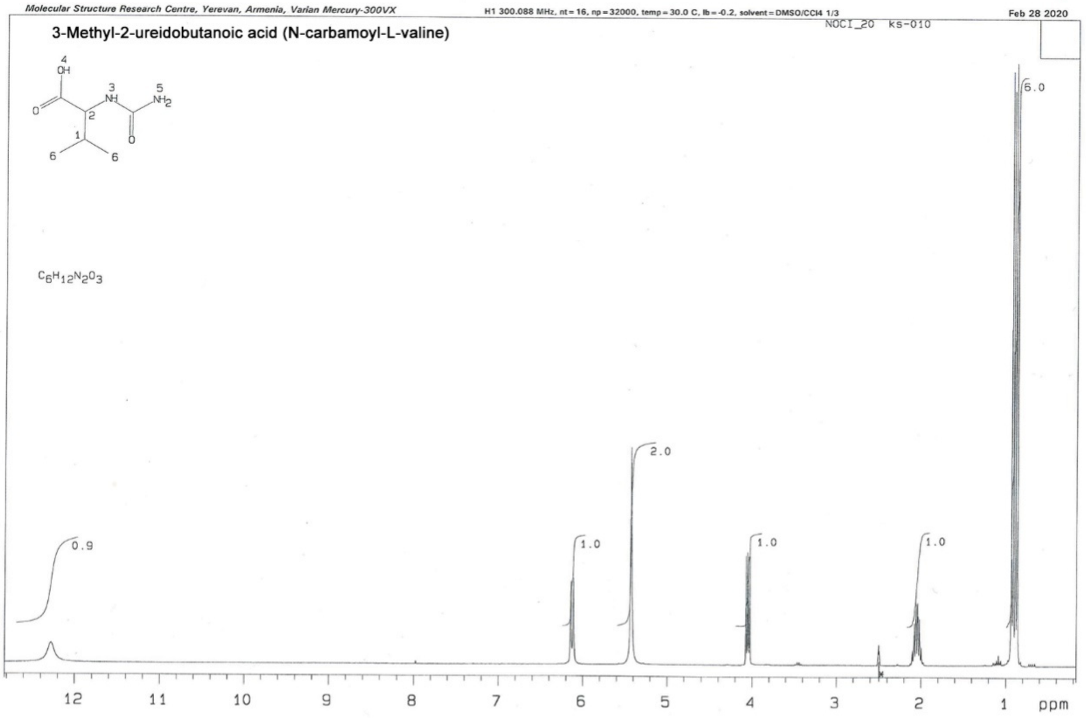


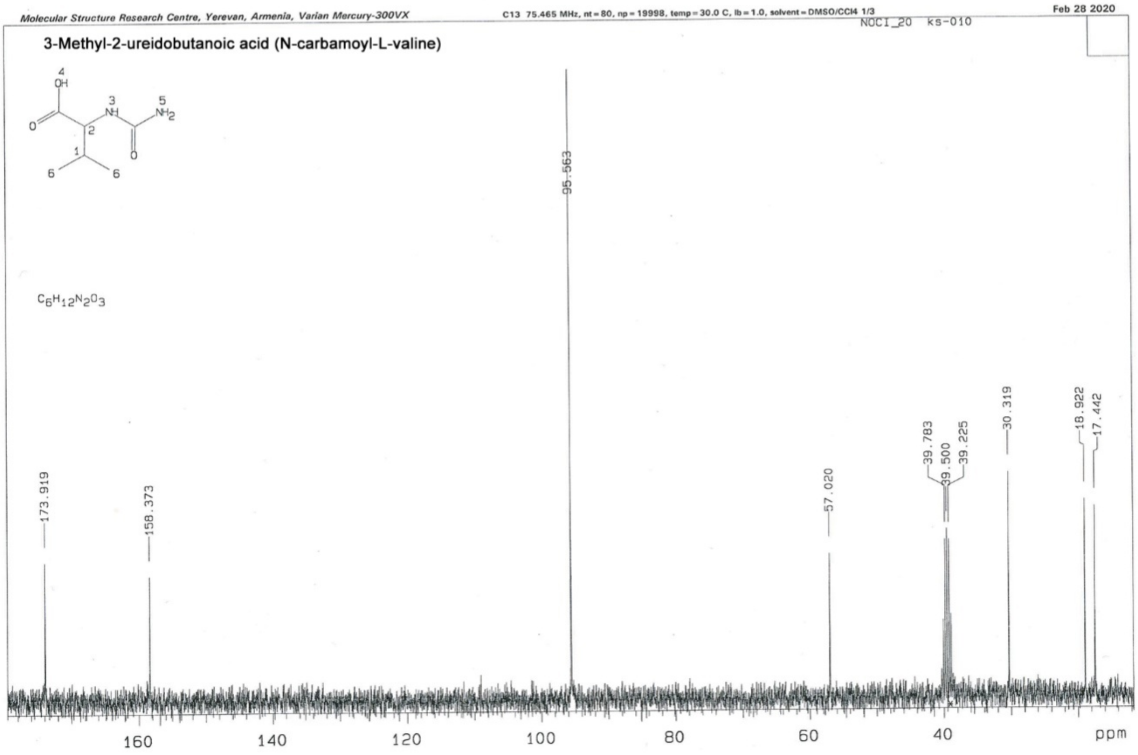

**Compound 11: 2-Ureidopropanoic acid.** From DL-Alanine. M. p. 187-189 °C (EtOH-$H_2O$, 2:1), lit.: 185 °C [4]. $^1H$ NMR δ: 1.28 (d, 3H, J = 7.3 Hz, $CH_3$); 4.11 (dq, 1H, J = 7.8, 7.2 Hz, CH); 5.41 (br. 2H, $NH_2$); 6.18 (br. d, 1H, J = 7.8 Hz, NH); 12.31 (br., 1H, COOH). $^{13}C$ NMR δ: 18.3 ($CH_3$); 47.7 (CH); 158.0 (NCO); 174.9 (OCO).

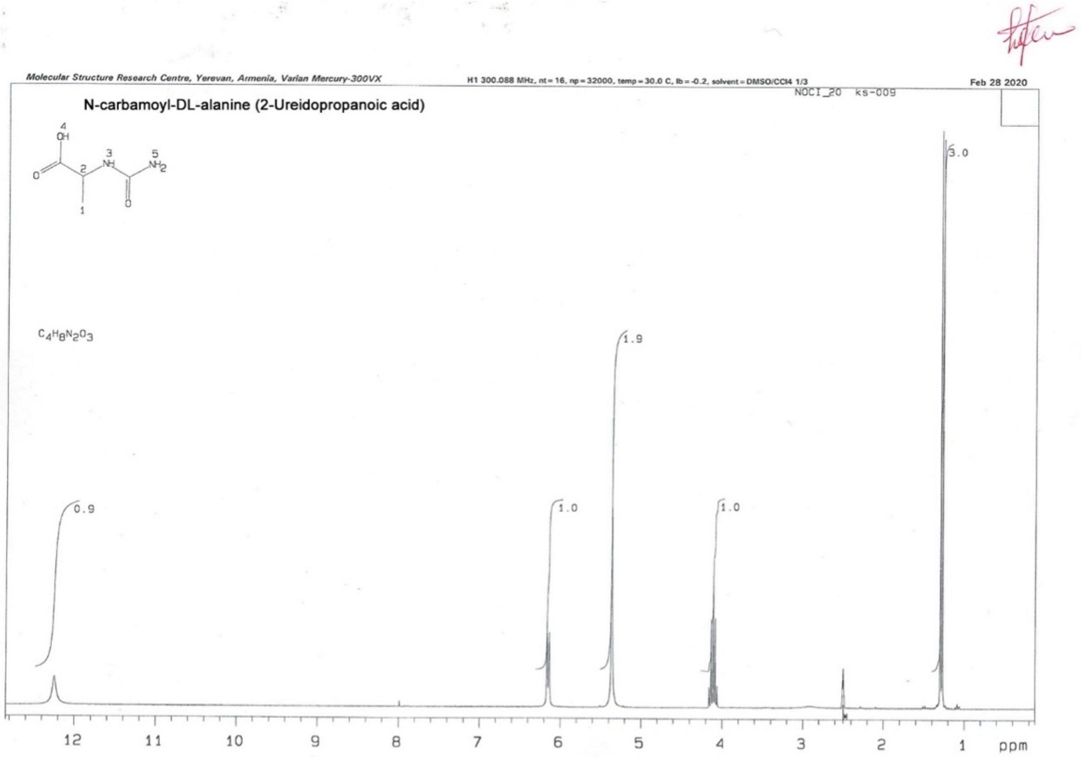


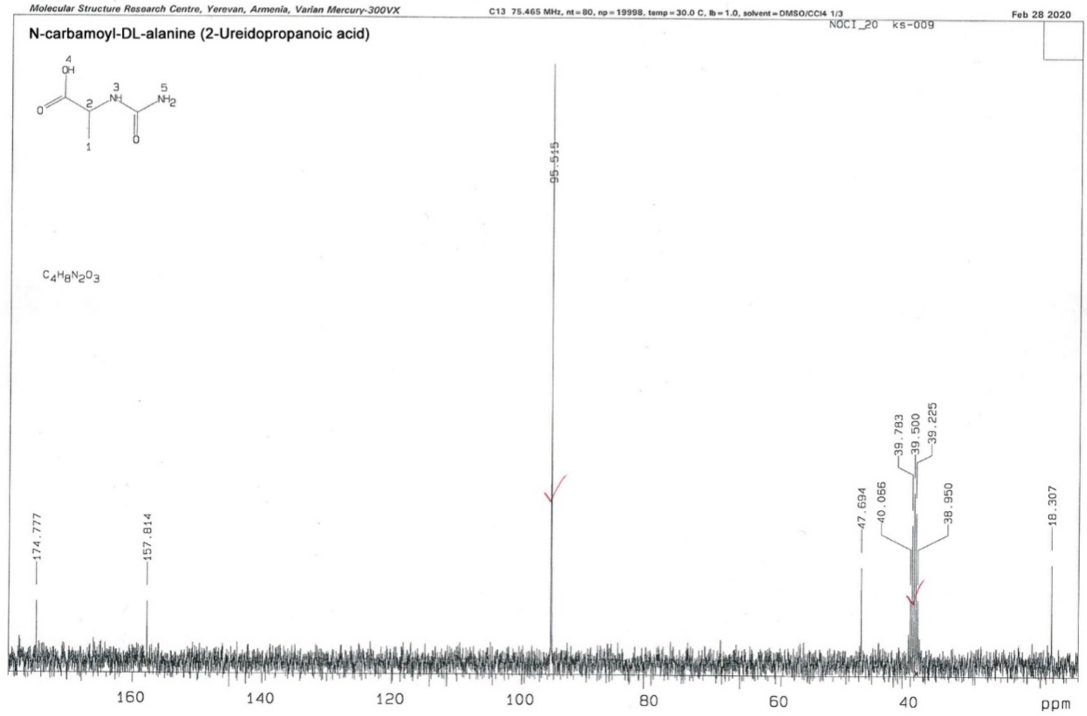

**Compound 12: 2-Ureidopropanoic acid.** From D-alanine. M. p. 201.5-202 °C (EtOH-$H_2O$, 2:1). Data of $^{1}H$ and $^{13}C$ NMR are same as for **11**.

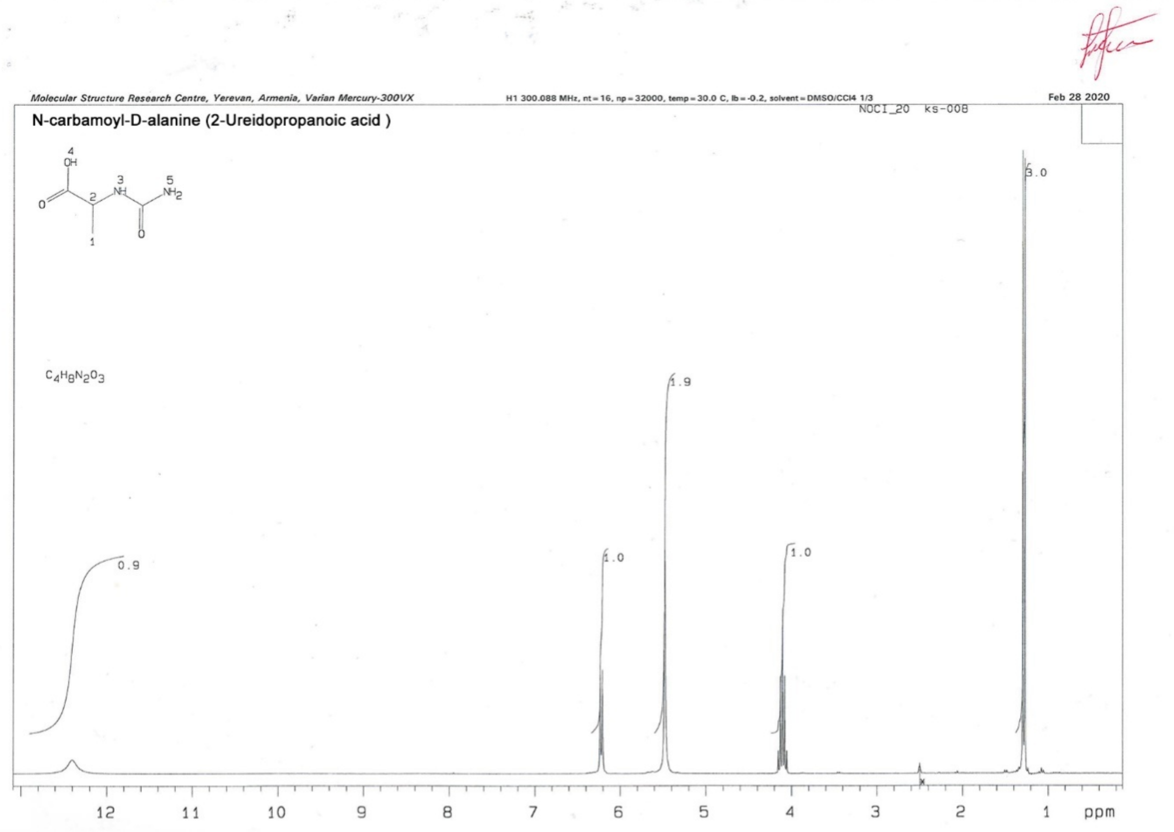


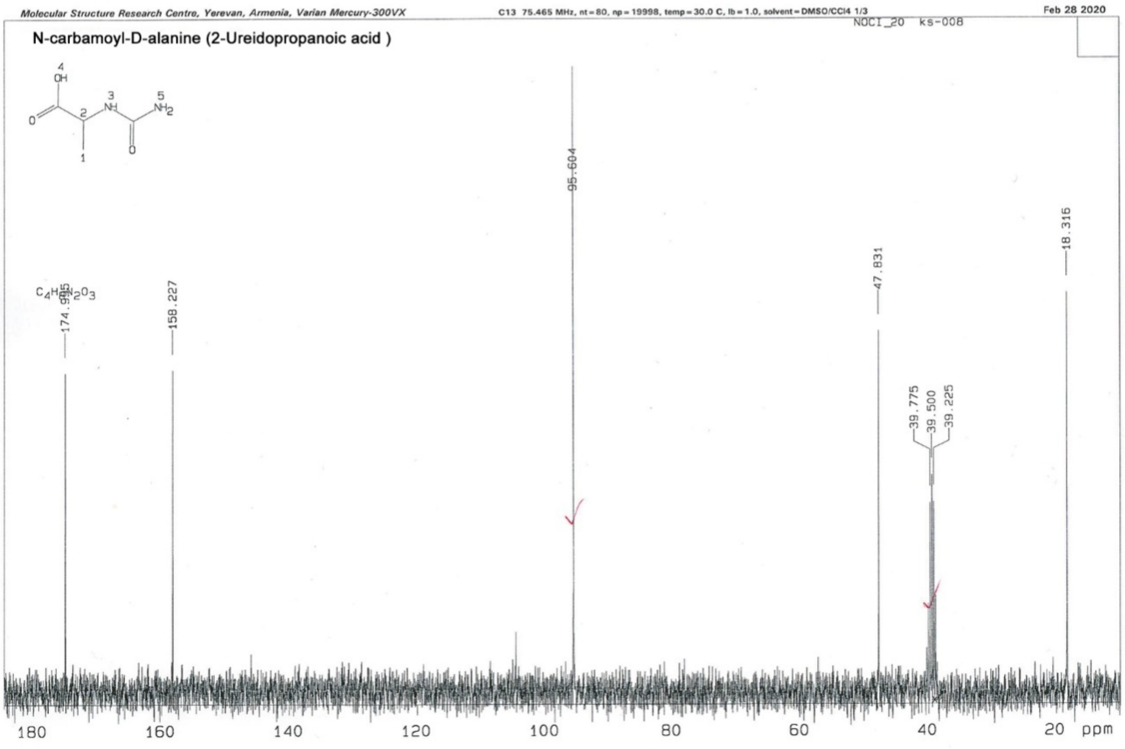

**Compound 13: 2-Ureidopropanoic acid.** From L-alanine. M. p. 208-209 °C (EtOH-$H_2O$, 2:1), lit.: 165-167 °C [1], 198-200 °C[4 ].Data of [1]H and [13]C-NMR are same as for **11**.

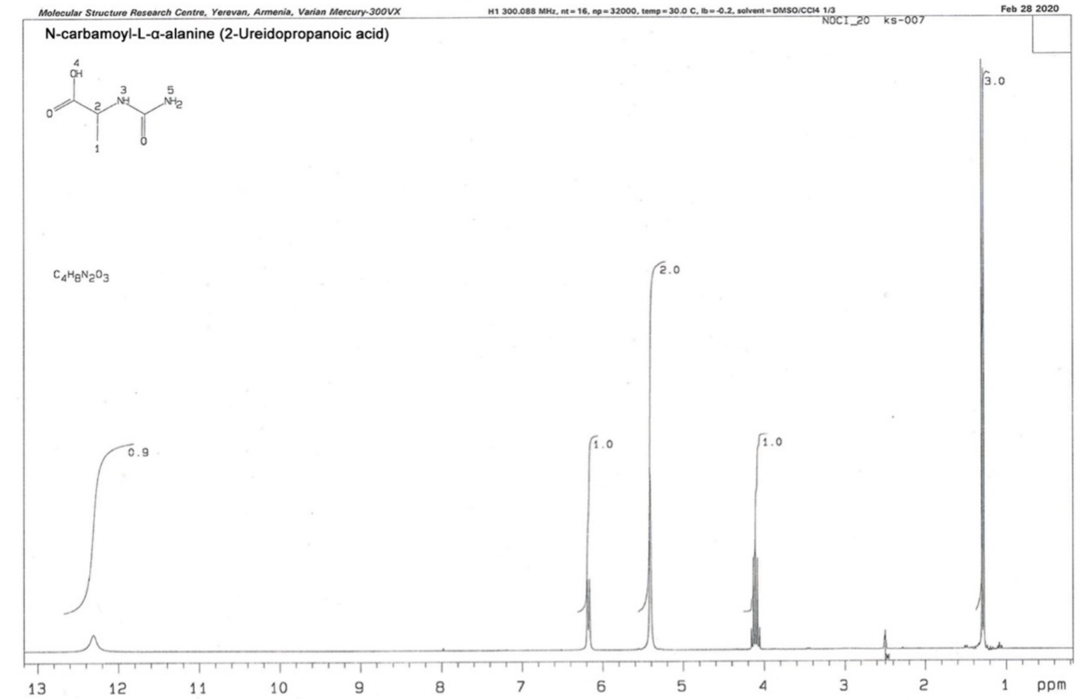


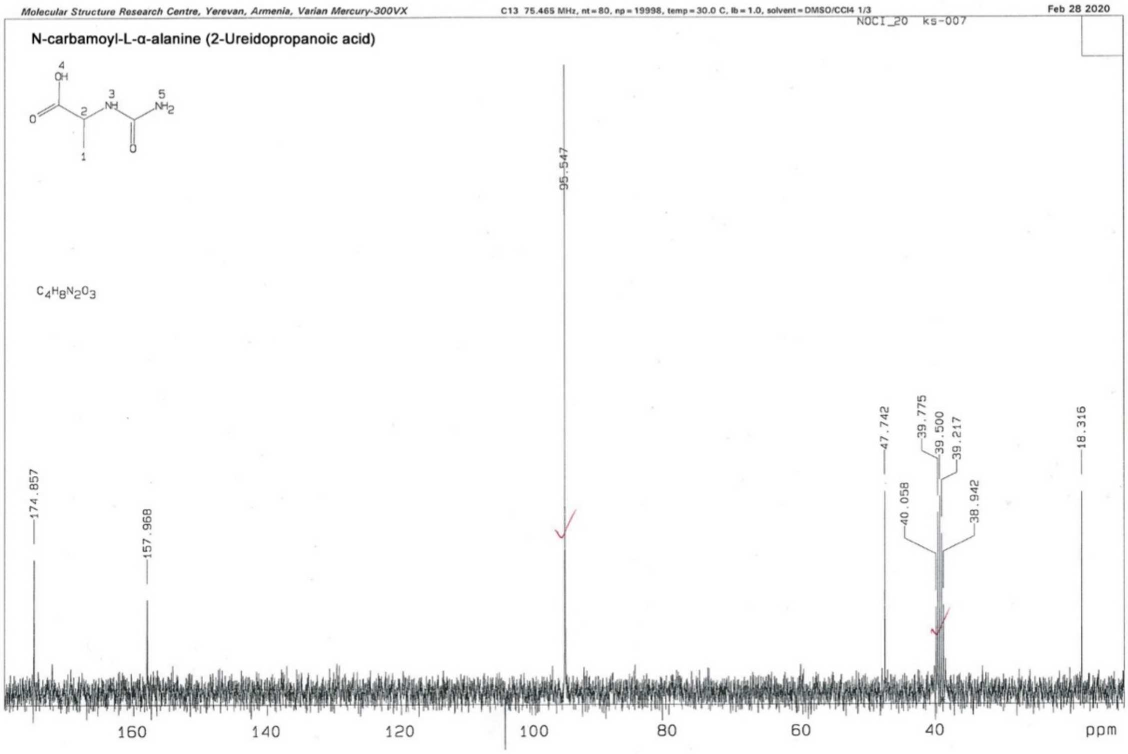

**Compound 14: N-Carbamoyl-3-(4-isopropoxyphenyl)-4-aminobutanoic acid.** From previously synthesized 3- (4-isopropoxyphenyl)-4-aminobutanoic acid [5].M.p. 217-219 ºC (EtOH).$^{1}$H NMR δ: 1.30 (d, 6H, J = 6.0 Hz, $2CH_3$); 2.38 (dd, 1H, J = 15.6, 8.4 Hz, $CH_2$); 2.59 (dd, 1H, J = 15.6, 5.7 Hz, $CH_2$); 3.04 – 3.28 (m, 3H, $NCH_2CH$); 4.51 (sp, 1H, J= 6.0 Hz, OCH); 5.21 (br., 2H, $NH_2$); 5.79 (br.t, 1H, J = 5.5 Hz, NH); 6.74 – 6.79 (m, 2H, Ar); 7.08 – 7.13 (m, 2H, Ar); 11,79 v.br.(1H, COOH). $^{13}$C NMR δ: 21.7 ($2CH_3$); 37,9 ($CH_2$); 41,3 (CH); 44,5 ($NCH_2$); 68.7 (OCH); 115.0 (2CH); 128.2 (2CH); 134.0; 155.8; 158.4; 172,9.

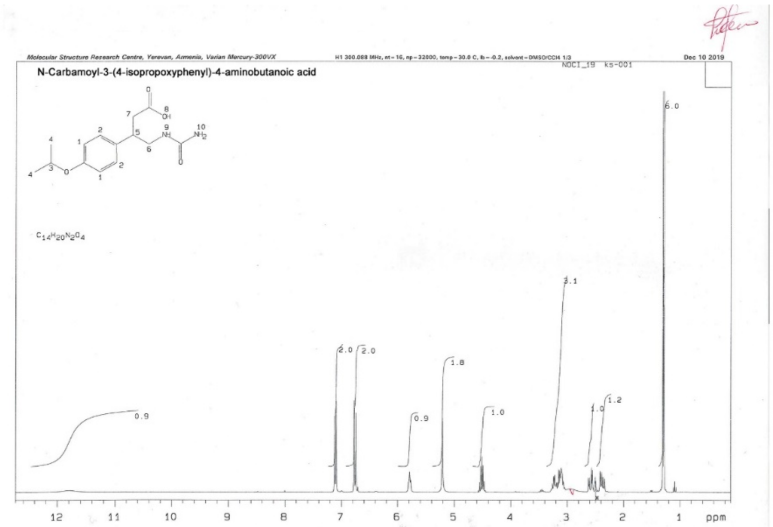


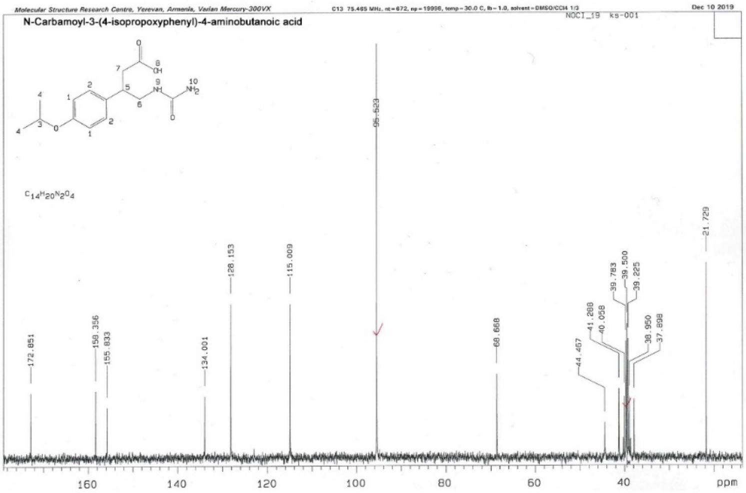


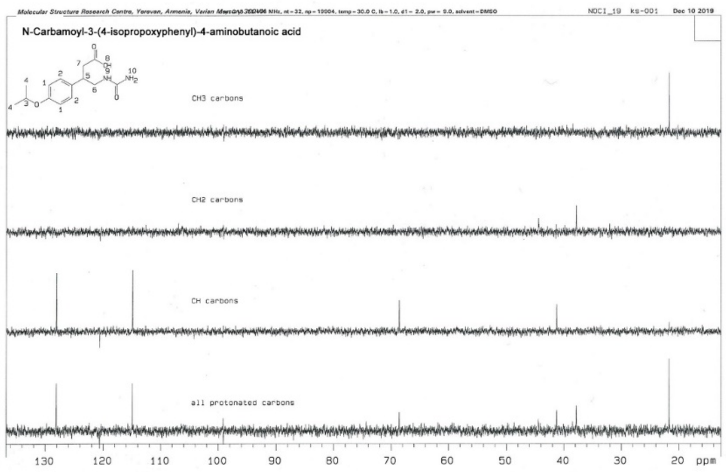

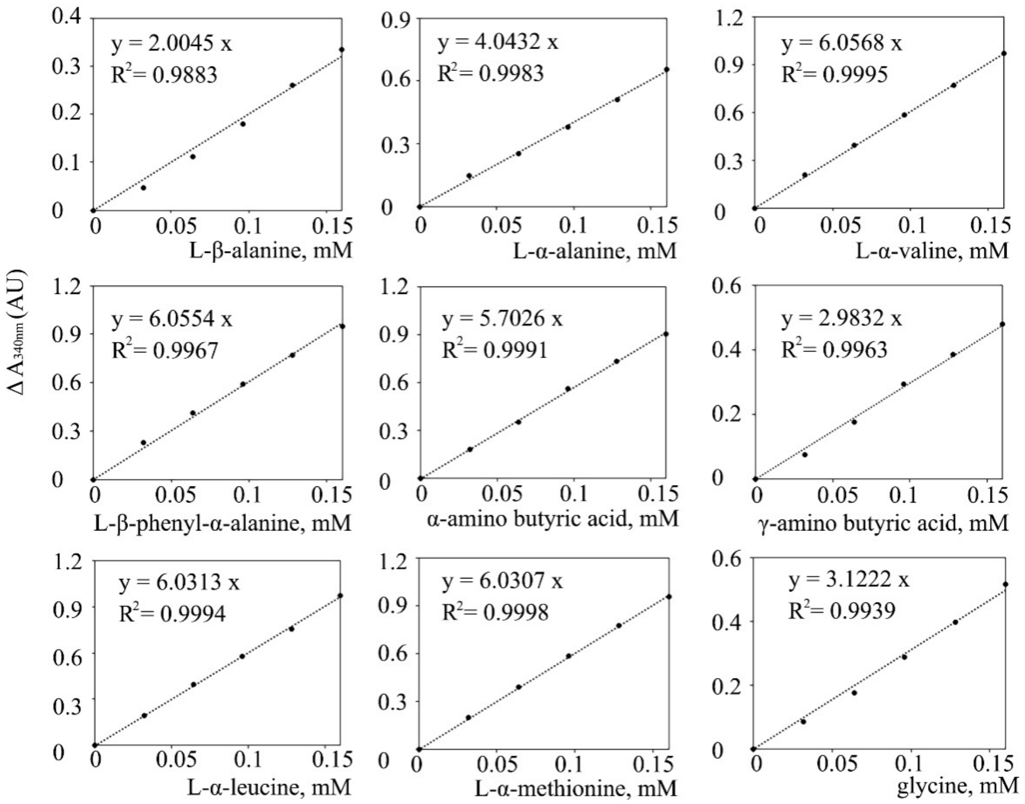


**Supplementary Figure 7. OPA derivatised amino acid standard curves for quantification of millimolar extinction coefficients.** All standard curves were generated by using a plot of absorbance vs. concentration at 340 nm to determine the molar extinction coefficient of amino acids. The samples were prepared at concentrations of 0, 0.032, 0.064, 0.096, 0.128 and 0.16 mM with 3 mL of freshly prepared activity reagent (0.1 M sodium borate pH 9.2, 2 mM OPA and 5 mM β-mercaptoethanol). After incubation at 20 °C for 30 min the absorption of the corresponding isoindole were determined at 340 nm. The fit lines were obtained by linear regression with $R^2$ greater than 0.988.

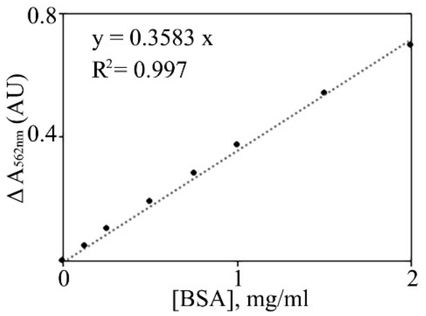


**Supplementary Figure 8. Bovine serum albumin standard curves for protein quantification by BCA.** BCA colour response curves produced using the Standard Test Tube Protocol (PierceTM BCA Protein Assay Kit) to quantify purified RrCβAA protein concentration. Raw data available at: https://doi.org/10.6084/m9.figshare.12399611.v1

#### Supplementary Tables

**Supplementary Table 1. Purification table of recombinant RrCβAA**

| Purification step | Protein concentration (mg/mL) | Specific activity (U/mg) | Yield (%) |
|---|---|---|---|
| Crude extract | 23.8 | 0.3 | 100.0 |
| His tag and concentration | 15.2 | 8.8 | 77.5 |
| Gel filtration and concentration | 12.6 | 13.4 | 65.2 |

**Supplementary Table 2. Temperature - activity relationship for RrCβAA**

| Temperature (°C) | Specific activity (U/mg) | Relative activity (%) |
|---|---|---|
| 25 | 2.7 ± 0.8 | 26±6 |
| 30 | 3.3±0.7 | 34±13 |
| 35 | 4.5±0.3 | 45±11 |
| 40 | 6.7±1.4 | 65±2 |
| 45 | 8.1±1.6 | 78±1 |
| 50 | 9.1±1.4 | 88±4 |
| 55 | 10.3±2.1 | 100±0 |
| 60 | 6.1±1.2 | 60±5 |
| 65 | 2.6±1.2 | 25±9 |
| 70 | 1.9±0.8 | 18±5 |

**Supplementary Table 3. Temperature - stability relationship for RrCβAA**

| Temperature (°C) | Specific activity (U/mg) | Relative activity (%) |
|---|---|---|
| 25 | 11.7 ± 0.5 | 100 ± 0 |
| 30 | 11.8 ± 0.4 | 101 ± 5 |
| 35 | 11.1 ± 0.4 | 95 ± 6 |
| 40 | 11.2 ± 0.4 | 96 ± 7 |
| 45 | 9.3 ± 0.8 | 80 ± 4 |
| 50 | 8.4 ± 0.7 | 71 ± 6 |
| 55 | 5.5 ± 0.9 | 47 ± 7 |
| 60 | 5.3 ± 0.7 | 45 ± 4 |
| 65 | 2.4 ± 0.9 | 21 ± 8 |
| 70 | 0.8 ± 0.3 | 7 ± 2 |

**Supplementary Table 4. RrCβAA activity in the presence of divalent cations and reducing agents**

| Sample | Specific activity (U/mg) | Relative activity (%) |
|---|---|---|
| As prepared | 13.6 ± 1.0 | 100 ± 7 |
| $Na^{+}$ | 13.4 ± 0.7 | 99 ± 5 |
| $K^{+}$ | 17.9 ± 2.0 | 131 ± 15 |
| $Ca^{2+}$ | 18.3 ± 1.0 | 135 ± 8 |
| $Mg^{2+}$ | 18.3 ± 1.3 | 135 ± 10 |
| $Ba^{2+}$ | 13.0 ± 1.6 | 95 ± 12 |
| $Sn^{2+}$ | 14.5 ± 1.0 | 107 ± 7 |
| $Pb^{2+}$ | 13.3 ± 1.3 | 98 ± 10 |
| $Fe^{2+}$ | 15.7 ± 2.2 | 116 ± 16 |
| $Fe^{3+}$ | 14.1 ± 2.1 | 103 ± 15 |
| $Mn^{2+}$ | 21.1 ± 2.5 | 155 ±18 |
| $Ni^{2+}$ | 25.5 ± 3.1 | 188 ± 22 |
| $Co^{2+}$ | 29.6 ± 4.0 | 218 ± 29 |
| $Cd^{2+}$ | 28.6 ± 2.9 | 210 ± 21 |
| $Zn^{2+}$ | 9.5 ± 0.6 | 70 ± 5 |
| $Cu^{2+}$ | 6.7 ± 0.5 | 49 ± 4 |
| β-mercaptoethanol | 15.0 ± 1.0 | 110 ± 8 |
| DTT | 12.6 ± 0.8 | 93 ± 6 |
| DTNB | 4.2 ± 0.2 | 31 ± 2 |
| EDTA | 0.7 ± 0.3 | 5 ± 2 |

**Supplementary Table 5. Activity recovery of EDTA inactivated RrCβAA enzyme**

| Sample | Specific activity (U/mg) | Relative activity (%) |
|---|---|---|
| As prepared | 13.1 | 100 |
| EDTA inactivated | 0 ± 0 | 0 ± 0 |
| $Na^{+}$ reactivated | 0.6 ± 0.1 | 4 ± 0 |
| $K^{+}$ reactivated | 0.1 ± 0.1 | 1 ± 0 |
| $Ca^{2+}$ reactivated | 1.1 ± 0.1 | 8 ± 1 |
| $Mg^{2+}$ reactivated | 1.4 ± 0.1 | 11 ± 0 |
| $Ba^{2+}$ reactivated | 0 ± 0 | 0 ± 0 |
| $Sn^{2+}$ reactivated | 0 ± 0 | 0 ± 0 |
| $Pb^{2+}$ reactivated | 0.3 ± 0.1 | 3 ± 0 |
| $Fe^{2+}$ reactivated | 1.6 ± 0.1 | 13 ± 0 |
| $Fe^{3+}$ reactivated | 1.9 ± 0.1 | 15 ± 1 |
| $Mn^{2+}$ reactivated | 41.5 ± 4.5 | 319 ± 38 |
| $Ni^{2+}$ reactivated | 42.4 ± 3.1 | 325 ± 24 |
| $Co^{2+}$ reactivated | 54.7 ± 3.1 | 418 ± 24 |
| $Cd^{2+}$ reactivated | 72.2 ± 4.6 | 552 ± 35 |
| $Zn^{2+}$ reactivated | 0 ± 0 | 0 ± 0 |
| $Cu^{2+}$ reactivated | 0 ± 0 | 0 ± 0 |

**Supplementary Table 6. Substrate preference of RrCβAA enzyme**

| Substrate | Specific activity (U/mg) |
| --- | --- |
| N-carbamoyl-L-β-alanine | 11.4 ± 3.4 |
| N-carbamoyl-L-α-alanine | 6.1 ± 1.8 |
| N-carbamoyl-D-alanine | 0.0 ± 0.0 |
| N-carbamoyl-DL-alanine | 6.3 ± 1.1 |
| N-carbamoyl-L-valine | 0.0 ± 0.0 |
| N-carbamoyl-D-valine | 0.0 ± 0.0 |
| N-carbamoyl-DL-valine | 0.0 ± 0.0 |
| N-carbamoyl-L- β-phenyl-α-alanine | 0.2 ± 0.1 |
| N-carbamoyl-L-β-phenyl- β -alanine | 0.0 ± 0.0 |
| N-carbamoyl-D-β-phenyl-α-alanine | 0.0 ± 0.0 |
| N-carbamoyl-DL-β-phenyl-α-alanine | 0.3 ± 0.1 |
| N-carbamoyl-α-amino butyric acid | 5.5 ± 1.1 |
| N-carbamoyl-γ- amino butyric acid | 7.9 ± 1.0 |
| N-carbamoyl-L-leucine | 0.2 ± 0.1 |
| N-carbamoyl-L-methionine | 4.6 ± 1.1 |
| N-carbamoyl-glycine | 8.4 ± 2.1 |
| N-carbamoyl-3-(4-isopropoxyphenyl)-4-aminobutanoic acid | 0.0 ± 0.0 |

**Supplementary Table 7. X-ray crystallography data collection and refinement statistics**

| | |
|---|---|
| **Data collection and analysis statistics** | |
| Beamline | DLS I04 |
| Date | 15/12/2019 |
| Wavelength (Å) | 0.912 |
| Resolution (Å) | 84.92 – 2.00 (2.05 – 2.00) |
| Space group | $P2_122_1$ |
| *Unit-cell parameters* | |
| a (Å) | 53.23 |
| b (Å) | 104.62 |
| c (Å) | 145.37 |
| α = β = γ (°) | 90.00 |
| Unit-cell volume (Å$^3$) | 809598 |
| Solvent content (%) | 46 |
| Measured reflections | 391722 (29077) |
| Independent reflections | 55817 (4087) |
| Completeness (%) | 100.0 (100.0) |
| Redundancy | 7.0 (7.1) |
| $CC_{1/2}$ (%) | 0.996 (0.786) |
| $\langle I \rangle / \langle \sigma(I) \rangle$ | 7.1 (1.3) |
| **Model refinement** | |
| Rwork (%) | 22.87 |
| Rfree (%) | 26.98 |
| No. of non-H atoms | |
| Protein | 6220 |
| Solvent | 362 |
| Ions | 4 |
| RMS deviation from ideal values | |
| Bond angles (°) | 1.85 |
| Bond lengths (Å) | 0.013 |
| Average B factors (Å$^2$) | |
| Protein | 31.7 |
| Solvent | 32.0 |
| Ions | 35.0 |
| Ramachandran plot | |
| Most favoured regions (%) | 98 |
| PDBID | 8C46 |

*Values for highest resolution shell are shown in parentheses.
